# Crowdsourcing Temporal Transcriptomic Coronavirus Host Infection Data: resources, guide, and novel insights

**DOI:** 10.1101/2022.12.14.520483

**Authors:** James Flynn, Mehdi M. Ahmadi, Chase T. McFarland, Michael D. Kubal, Mark A. Taylor, Zhang Cheng, Enrique C. Torchia, Michael G. Edwards

## Abstract

The emergence of SARS-CoV-2 reawakened the need to rapidly understand the molecular etiologies, pandemic potential, and prospective treatments of infectious agents. The lack of existing data on SARS-CoV-2 hampered early attempts to treat severe forms of COVID-19 during the pandemic. This study coupled existing transcriptomic data from SARS-CoV-1 lung infection animal studies with crowdsourcing statistical approaches to derive temporal meta-signatures of host responses during early viral accumulation and subsequent clearance stages. Unsupervised and supervised machine learning approaches identified top dysregulated genes and potential biomarkers (e.g., CXCL10, BEX2, and ADM). Temporal meta-signatures revealed distinct gene expression programs with biological implications to a series of host responses underlying sustained Cxcl10 expression and Stat signaling. Cell cycle switched from G1/G0 phase genes, early in infection, to a G2/M gene signature during late infection that correlated with the enrichment of DNA Damage Response and Repair genes. The SARS-CoV-1 meta-signatures were shown to closely emulate human SARS-CoV-2 host responses from emerging RNAseq, single cell and proteomics data with early monocyte-macrophage activation followed by lymphocyte proliferation. The circulatory hormone adrenomedullin was observed as maximally elevated in elderly patients that died from COVID-19. Stage-specific correlations to compounds with potential to treat COVID-19 and future coronavirus infections were in part validated by a subset of twenty-four that are in clinical trials to treat COVID-19. This study represents a roadmap to leverage existing data in the public domain to derive novel molecular and biological insights and potential treatments to emerging human pathogens. The data from this study is available in an interactive portal (http://18.222.95.219:8047).

## Introduction

The respiratory disease known as COVID-19 is caused by the *Severe acute respiratory syndrome-related coronavirus 2* (SARS-CoV-2), a species individuum within the *Betacoronavirus* genus and *Coronaviridae* family (Gorbalenya et al., 2020). Since emerging as a deadly human pathogen in December 2019, it has triggered a devastating global pandemic that may pervade humanity for years to come (Shereen et al., 2020). While SARS-CoV-2 is the causative agent of COVID-19, it is just one of many respiratory viruses that arise and reemerge regularly. Vaccination has been shown to mitigate severe responses to SARS-CoV-2, but as the best therapeutic option to date, it does not always prevent infection nor hospitalization (Excler et al., 2021, Koff et al., 2021, Lopez Bernal et al., 2021). As evident in this pandemic, there is need for more therapies that can act before new vaccines are developed to emerging pathogens, treat unvaccinated people and those infected by vaccine resistant variants (Hoffmann et al., 2020).

Many aspects of the cellular and molecular biology of the host infection response to SARS-CoV-2 are not well understood given its recent presumed zoonotic emergence as a human pathogen. The SARS-CoV-2 single positive strand RNA genome shares a 79% nucleotide identity to SARS-CoV-1, which caused a much smaller pandemic in 2002 (Abdelrahman et al., 2020, Lu et al., 2020, Wu Aiping et al., 2020, Zhou et al., 2020). Both viruses appear to have originated in bats and bind the Angiotensin Receptor 2 (ACE2) on host cells mainly via the viral spike protein to initiate cellular entry (Wan et al., 2020). The *Middle East respiratory syndrome-related coronavirus* (MERS-CoV) caused international outbreaks in 2012 and 2015 (Lu et al., 2020, Zaki et al., 2012); it differs from SARS-CoV viruses by binding to the Dipeptidyl Peptidase 4 (DPP4) receptor for cell entry (Li et al., 2019, Raj et al., 2013). A more complete understanding of coronavirus host infection mechanisms is necessary in preparation for combatting future outbreaks.

Coronavirus genomes are unusually large compared to other viruses (Masters, 2006). The 26,000 to 32,000 nucleotide genomes contain a varying number of open reading frames that encode 20 or more proteins, including a set of highly conserved replicase enzymes and the nucleocapsid, spike, membrane and envelope structural proteins in SARS-CoV-1, SARS-CoV-2 and MERS-CoV (Li et al., 2019, Song et al., 2019, Wu Aiping et al., 2020). Less conserved accessory proteins localize broadly to the cell surface, cytoplasm, various organelles, and the nucleus of host infected cells (Tan et al., 2006). The breadth of functional effects is impressive. A deeper understanding of common viral induced adverse cellular events and the temporal order of host molecular and pathway level response could provide opportunities for therapeutic intervention irrespective of infecting *Betacoronavirus* type (Totura and Bavari, 2019).

Meta-analysis workflows integrate observations across independent studies (Glass, 1976). More robust insights can be gained compared to single studies because each study has inherent variability that can be minimized when assessing a set of related studies. Variability embedded in medical research can originate, for example, in the methods of sample collection, experimental platform (e.g., microarrays *vs.* sequencing) or in the pipeline of algorithms used to process raw data. Meta-analysis can be done by merging datasets together or by using ensemble methods that utilize “wisdom of crowds” approaches (Galton, 1907, Sparks et al., 2016). The latter crowdsourcing approach has distinctive advantages as overall error in the “crowd” or collection of datasets can be minimized when factoring each individual study error. By using the same comparison criteria (e.g., coronavirus infected lung tissue *vs.* mock infected lung tissue), crowdsourcing results can faithfully build observational consensus and reliability.

The current study applied a novel “wisdom of crowds” technique to generate temporal gene expression meta-signatures from 15 independent RNA expression studies comprising 74 SARS-CoV-1 lung infected *vs.* mock infected comparisons in mouse at 0.5, 1, 2, 4, and 7 days post-infection. The meta-signatures provide a robust molecular representation of disease progression that has been supported by emerging studies on SAR-CoV-2. Furthermore, functionally inverse correlations to mechanistically distinct compound treatments provide a means to anticipate potential benefit of single or combination treatments.

The overarching goals of this study are to: i) present a general guide to crowdsourcing publicly available transcriptomic data, ii) provide a resource of data and metadata on SARS-CoV infection, iii) identify drugs with potential to treat specific biomarker-based stages of disease and iv) derive actionable insights of SARS-COV-2 host response in humans. The authors invite engagement of the coronavirus research community by providing data, tools, and resources to help advance research on coronavirus infectivity. An interactive portal is available as a public resource (http://18.222.95.219:8047).

## Methods

Illumina’s BaseSpace™ Correlation Engine (Kupershmidt et al., 2010) (BSCE, previously known as NextBio) was used to facilitate the aggregation and mining of coronavirus host infection transcriptome data for two main reasons. BSCE contains over 21,500 omic-scale Curated Studies with over 132,000 experimental comparison results (hereafter called biosets). Consistent curation practices ensure the inclusion of high quality datasets for analysis and improves the resulting interpretations. Curated Studies in BSCE are processed in standard, platform-specific pipelines to generate gene sets and measures. For each study, available Test group *vs.* Control group results plus experimental metadata are generated. Second, BSCE has robust data-driven correlation, aggregation, and machine learning applications to exploit the consistent processing and curation of omic-scale studies derived from international public repositories, including US-NCBI’s Gene Expression Omnibus (Barrett et al., 2013) and the Short Read Archive (Leinonen et al., 2011), EMBL-EBI’s Array Express (Athar et al., 2019), US-NIEHS’s NTP Chemical Effects in Biological Systems (CEBS) (Gwinn et al., 2020), and Japan’s Toxicogenomics Project-Genomics Assisted Toxicity Evaluation System (TG-GATES) (Igarashi et al., 2015).

### Systematic Review of Omic-scale Studies and Experimental Designs

A search of BSCE identified 44 Curated Studies with a total of 447 biosets querying the term “Coronavirus”. The biosets covered a wide range of experimental designs across an array of viral species, viral individuum variants, viral dose, post-infection duration, host organism, host tissue, host sex, host age, gene knockout models, cell line models, compound treatments and technology platforms. Study and experimental bioset details were hand curated from BSCE, the GEO entry and primary articles.

### RNA Expression Analysis

Threshold filters between test (e.g., virus infected) and control (e.g., mock infection) conditions were applied to log2 scale RNA expression data. Genes with both mean normalized test and control intensities less than the 20^th^ percentile of the combined normalized signal intensities were removed. A parametric Welch t-test, where variances are not assumed equal, is calculated and genes with a p-value > 0.05 were removed. An additional filter removes genes with non-log2 absolute fold change < 1.2 to generate the final list of differentially expressed genes in a bioset. The parametrically determined gene list is set to non-parametric ranks by assigning the gene with the highest absolute fold change the highest rank of 1 and genes descending by absolute fold change are numbered down the ordinal ranks. Bioset-Bioset correlations are done based on the ranks using the Running Fisher algorithm (Kupershmidt et al., 2010). BSCE calculates normalized, bioset-specific gene scores that considers the absolute fold change rank, the number of genes passing threshold filters and the number of genes measured by the platform on a 100 to 0 scale. A positive or negative sign was appended to gene scores based on the fold change direction to generate values ranging from -100 to +100, here forth called directional gene score(s).

Initial RNA expression analysis of SARS-CoV-1 lung infected *vs.* mock infected control used biosets from 4 mammalian species and 9 beta-coronavirus variants, encompassing 19 Curated Studies with a total of 97 biosets [*Mus musculus* - mouse (n=85), *Macaca fascicularis* - cynomolgus macaques (n=5), *Mustela putorius furo* - ferret (n=5) and in *Homo sapiens* - human cells (n=2)]. Biosets covered 48 early, 42 mid and 7 late stage infection results [0.5, 1, 2, 3, 4, 5, 7, 14, 21 and 28 days post-infection (dpi)]. Beta-coronavirus variants included SARS-CoV-MA15, SARS-CoV-MA15e, SARS-CoV-MA15g, SARS-CoV-TOR-2, SARS-CoV-GZ02, SARS-CoV-HC-SZ-6103, SARS-CoV-ic, SARS-CoV-Urbani, and SARS-CoV-2 USA-WA1/2020 (NR-52281). A lethal SARS-CoV-MA15 virus was created by serial passage of the natural SARS-CoV-Urbani variant in the lungs of mice (Roberts et al., 2007). Technology Platforms included [Agilent-014868 Whole Mouse Genome Microarray 4x44K G4122F, Affymetrix Mouse Genome 430 2.0, Affymetrix Canine Genome 2.0 Array, Array Affymetrix Rhesus Macaque Genome Array, Illumina NextSeq 500]. We filtered the time course based on the number of biosets available per dpi; those with less than 3 biosets were excluded to minimize skewing from an overreaching but underrepresented variable set. This left 74 SARS-CoV-1 lung infected *vs.* mock infected biosets in mouse at 0.5, 1, 2, 4, and 7 dpi comprised of 299 experimental and 239 control animals as the focus for the current study. Metadata and a gene expression matrix are in Supplemental Table 1.

### Generation and Analysis of Temporal Meta-Signatures

The 74 biosets spanned i) five post-infection time points, ii) 3 host age groups, iii) 3 host strains, iv) 5 viral variants, and v) 4 magnitudes of viral dose. Age groups were divided into young mice (<= 10 wks), adult (20 to 23 wks), and aged (52 wks). To the best of our knowledge all were female. *Mus musculus* strains included BALB/c, 129/S6/SvEv and C57BL/6NJ. The most commonly used MA15 viral variant induces more severe reactions in mice than MA15g, MA15e or the ic (infectious clone) and wildtype TOR-2 variants (Roberts et al., 2007, Rockx et al., 2009). Intranasal infection dose levels included 100, 1,000, 10,000, 22,000 and 100,000 viral plaque forming units (pfu). As with humans afflicted with SARS-CoV-2, younger hosts appear less susceptible to the effects of the SARS-CoV-MA15 virus than aged mice (Frieman et al., 2010). Severity across age groups was also measured by the number of genes differentially expressed as viral inoculation dose increased over four magnitudes.

The BSCE Meta-Analysis application takes an input of a collection of biosets and outputs a table with the top 5,000 genes ranked in rows by highest sum of the normalized gene scores with bioset-specific fold change and p-values in columns. Meta-Analysis was run separately in sets for 6 biosets at 0.5 dpi, 15 biosets at 1 dpi, 19 biosets at 2 dpi, 19 biosets at 4 dpi and 15 biosets at 7 dpi. The average of the directional gene score values was taken for each gene in the set, here forth called temporal gene score(s), to generate five temporal meta-signatures that were aligned into a matrix to form 10,564 gene-specific time vectors following host response to SARS-CoV-1 infection in mouse lungs (Supplemental Table 2). The Venn diagram was generated using the Gent University Bioinformatics and Evolutionary Genomics application at http://bioinformatics.psb.ugent.be/webtools/Venn/. Profile charts were generated using JMP^®^ Pro 15.0 (SAS Institute Inc., Cary, NC).

The BSCE Pathway Enrichment application was used to correlate and rank the temporal meta-signatures to functional biogroup lists derived from Gene Ontology (GO) (Ashburner et al., 2000, Gene Ontology Resource, 2021). Biogroup concepts with a minimum of 10 and a maximum of 750 genes were used, thereby eliminating minor concepts and those with little specificity.

Network Analysis was performed using STRING, Protein-Protein Interaction Networks and Functional Enrichment Analysis at https://string-db.org (Szklarczyk et al., 2019) and Pathway Studio (Elsevier B.V., Amsterdam, Netherlands). Other gene function analyses were performed using the WEB-based GEne SeT AnaLysis Toolkit software (http://www.webgestalt.org/) (Liao et al., 2019, Wang et al., 2017).

### Unsupervised and Supervised Machine Learning Analyses

Principal Component Analysis of the 74 biosets used Partek Genomics Suite Version 7.21.1119 with PC product values, ellipsoids were graphically rendered at 2 standard deviations from centroid points. Partek was also used for F-value generation (between group variance / within group variance) to provide measures for experimental sources of variation. The F Distribution Calculator at https://www.statology.org/f-distribution-calculator/ was used to derive p-values using experimental variable degrees of freedom and the F-values.

Agglomerative hierarchical clustering for the 74 biosets and for the five temporal meta-signatures was performed separately by generating a static heatmap using the native R heatmap function “heatmap.2”, which orders features based on Euclidian distance and nearest neighbors agglomeration, no additional normalization was performed, and no parameters adjusted. Row and column dendrograms were isolated from the static heatmap and applied to third party heatmap packages “heatmaply” (Galili et al., 2018) and “InteractiveComplexHeatmap” (Gu and Hübschmann, 2021) to generate interactive heatmaps.

For supervised machine learning a binary elastic-net logistic regression combined with bootstrapping was applied to derive predictive models and robust coefficients of the genes in making temporal predictions (Barretina et al., 2012, Zou and Hastie, 2005). Given observations of distinctive early *vs.* late activities, samples were grouped as either early [0.5, 1, 2 dpi] or late [4, 7 dpi] to derive a small set of genes that could predict the stage of SARS-CoV-1 infection. Each bootstrap split the data into different training and testing sets with a ratio 4:1 using the “train_test_split” function from sklearn (Pedregosa et al., 2011) with “stratify=y” and a random seed generated using “randrange” function. One hundred different splits were generated by repeatedly calling “randrange” function in “train_test_split” and any duplicated splits were programmatically removed. For each split, a combination of 3-fold cross-validation and hyperparameters search was performed using the “GridSearchCV” function with “cv=3” from sklearn on the training set. The two hyperparameters, “C” and “l1_ratio” were the targets of the grid search. The “C” parameter is the inverse of lambda, which controls the overall strength of the regulation term in elastic net, and was set to 1e-4, 1e-3, 1e-2, 1e-1, 1 and 10. The “l1_ratio” controls the ratio between L1 and L2 regulation terms, which was 0.2, 0.4, 0.6, 0.8 and 1. The combination of these two hyperparameters with the lowest cross validation error was selected and used to derive the coefficient for each gene in the model to predict the early or late stage for each sample in the validation and test sets. The validation accuracy was reported as training accuracy and the test accuracy was calculated by using “accuracy_score” function from sklearn by using the best model on testing set. The coefficients of the genes from these 100 bootstraps were summarized and only genes with at least 90 of 100 bootstraps with a nonzero weight were reported in the final list with the average coefficient and standard deviation.

### Correlations to Compound Treatments, Gene Perturbations and Diseases

The five temporal meta-signatures were imported into BSCE as biosets and queried in the BSCE Pharmaco, Knockout, Knockdown and Disease Atlas applications. Bioset queries made against the entire repository yield a Bioset-Bioset correlation rank ordered list. BSCE biosets are tagged with concept terms that capture sample and experimental details, like compound treatments, genetic perturbations and phenotypes. A machine learning categorization process assesses the rank of correlations to the tagged concept term results into Pharmaco, Knockdown and Disease Atlases using relevant tag types and strength of contributing bioset correlations (BSCE Supplemental Text). The BSCE Pharmaco Atlas application was used to categorize, and rank compounds based on meta-signature correlations to all ‘treatment *vs.* control’ biosets in the system [4,804 Compounds in 53,723 biosets from 6,852 Studies]. The BSCE Knockdown Atlas application was used to categorize, and rank genes based on meta-signature correlations to all ‘gene perturbation *vs.* control’ biosets in the system [5,341 genes perturbed in 37,838 biosets from 10,287 Curated Studies]. The BSCE Disease Atlas application was used to categorize, and rank disease phenotypes based on queries with i) the genes *Cxcl10* and *Bex2* and ii) temporal meta-signature correlations to all ‘disease phenotype *vs.* normal’ biosets in the system [5,309 Phenotypes in 68,697 biosets from 12,929 Curated Studies].

Compound scores, and separately gene knockout scores, were scaled from -100 to +100 for each meta-signature timepoint and aggregated into a matrix of correlative perturbation time vectors over the course of SARS-CoV-1 infection. Compounds known to be in Clinical Trials were derived from https://ghddi-ailab.github.io/Targeting2019-nCoV/clinical

### Correlations to Human COVID-19 single cell RNAseq and Plasma Proteomic Studies

Single cell RNAseq data was obtained from (Liao et al., 2020), (Schulte-Schrepping et al., 2020), (Wilk et al., 2020), (Lee et al., 2020). Tables from each of the articles were downloaded and divided into individual cell type specific bioset files and imported into BSCE as Curated Studies for bioset-bioset correlations to temporal meta-signatures and gene clusters.

Plasma proteomic data generated from 384 patients presenting with acute respiratory distress in a hospital Emergency Department, 306 were COVID-19 positive (blood sampled on days 0, 3, 7, 28) and 78 non-COVID-19 controls (sampled only on day 0) was downloaded from https://www.olink.com/application/mgh-covid-19-study (Filbin et al., 2021). Disease severity was divided into 6 categories of increasing progression: 1) Not hospitalized, 2) Hospitalized, no supplemental oxygen required, 3) Hospitalized, supplemental oxygen required, 4) Hospitalized, Intubated, ventilated, and survived to 28 days, and 5) Death within 28 days. WHOmax was captured as the patient’s most severe category over time.

### Interactive Portal

The 74 Bioset and the 5 Temporal Meta-Signature results were collaboratively explored using Illumina Connected Analytics (https://www.illumina.com/products/by-type/informatics-products/connected-analytics.html) and then developed into an interactive portal using Dash by Plotly (http://conference.scipy.org/proceedings/scipy2019/shammamah_hossain.html) with the following features: i) Table View with gene search and meta-signature gene profile plots, ii) Principal Component Analysis, and iii) agglomerative clustering heatmaps. Access is available at http://18.222.95.219:8047. The data and code are also available at GitHub (https://github.com/Mark-A-Taylor/Coronavirus_Portal.git/).

## Results

The development of temporal meta-signatures of host coronavirus infection and the process used to derive biological interpretation is presented in Figure 1. The approach followed 8 main steps: 1) an assessment of available coronavirus Curated Studies in BSCE, 2) aggregating biosets of interest, 3) applying the BSCE Meta-Analysis application to provide normalized gene scores, 4) creation of meta-signatures, 5) functional enrichment, 6) network analysis 7) ranking of genes knocked out based on the strength of correlations to gene perturbation, and 8) ranking of compounds based on the strength of correlations to compound treatment.

**Figure 1.**
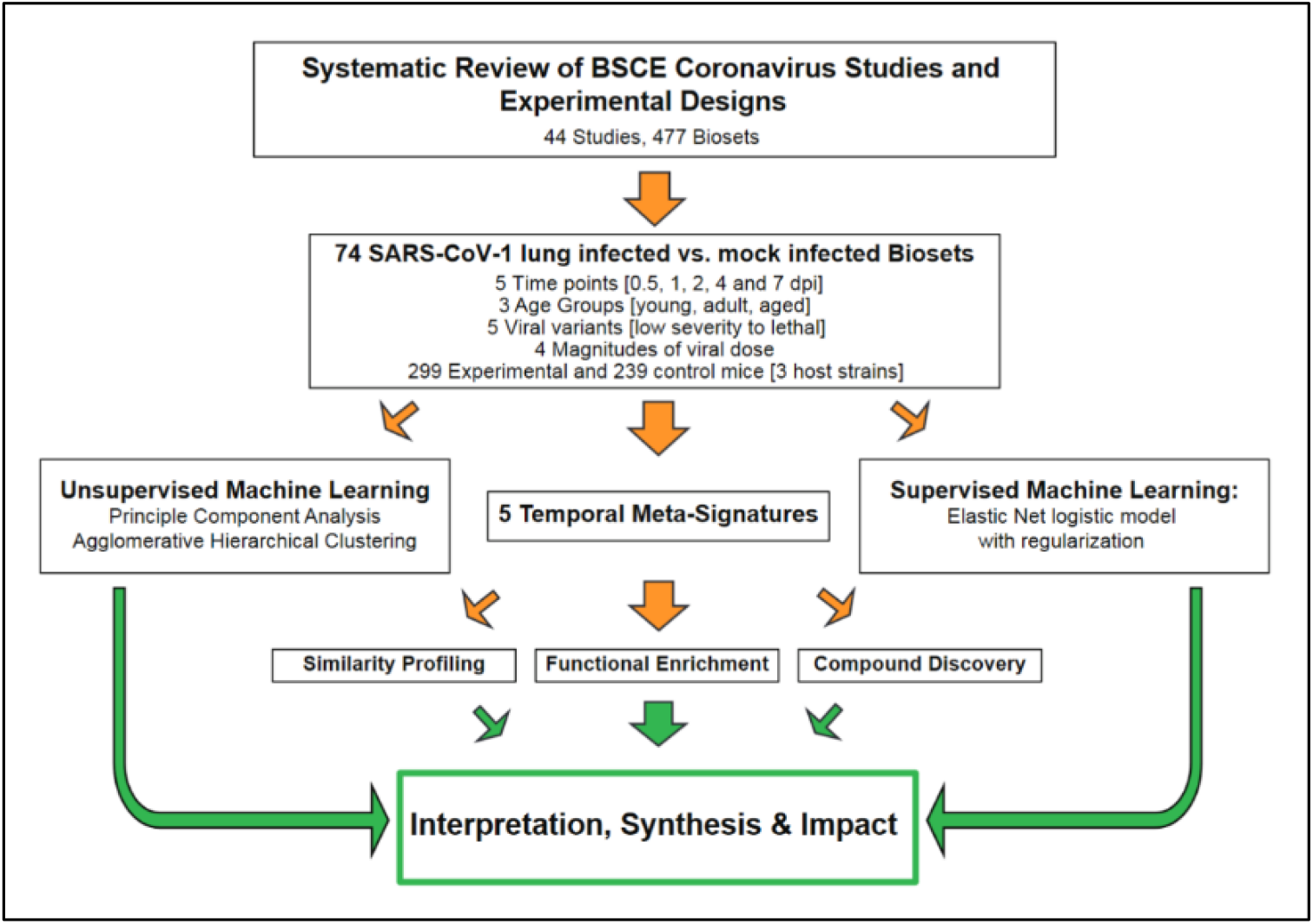
Workflow For Coronavirus Host Infection Biomarker Discovery. The schematic diagram outlines the process to derive and analyze an aggregate gene signature from an unbiased accounting of all coronavirus RNA Expression data available from Gene Expression Omnibus and in BaseSpace™ Correlation Engine (BSCE) Curated Studies. dpi = days post infection.

### I. Analysis of 74 SARS-CoV-1 Lung Infection Biosets

To combine data from different studies, datasets were normalized using fold change based gene score values on a scale of -100 to +100 (directional gene score) (see Methods and Supplemental Table 1). Figure 2A shows an example of the distributions of gene expression values from two biosets at 2 dpi before and after normalization. One bioset is from lungs of young mice infected with the less severe TOR-2 coronavirus variant *vs.* mock infected controls run on an Affymetrix microarray platform; the other is from lungs of aged mice infected with a lethal MA15 coronavirus variant *vs.* mock infected controls run on an Agilent microarray platform. The graphs show the normalized datapoints have a better linear fit than unnormalized fold change values (R^2^ of 0.683 *vs.* 0.507, respectively), are more evenly distributed and better suited for ensemble meta-analysis.

**Figure 2.**
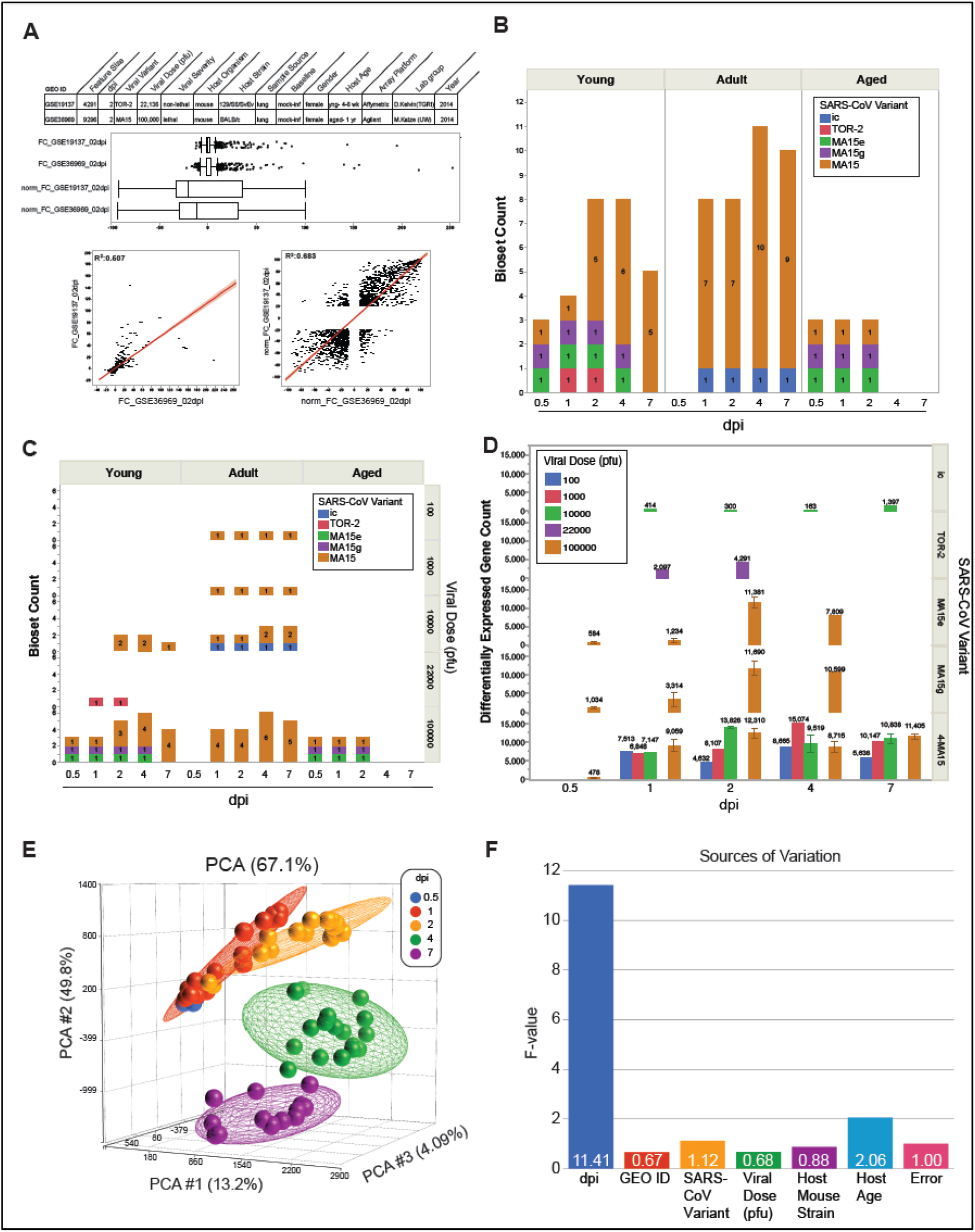
Coronavirus Bioset Normalization and Attribute Distribution. (A) Graphs show the decisive effects of directional gene score normalization *vs.* fold change values for 2 biosets compared across different model systems and technologies at 2 dpi. The top panel shows box plots with the distribution of the directional gene scores *vs.* fold changes. Rank-based scoring of fold change data converts biosets into a uniform -100 to +100 directional gene score list that accounts for a platforms’ number of measured genes and the feature size of each bioset. (B) Bioset count breakdown by age group, dpi and viral variant (C) Bioset count breakdown by age group, dpi. viral variant and viral dose. (D) Differentially expressed gene count breakdown by dpi, viral variant and viral dose. +/- SE shown where applicable. (F) Sources of variation plot across experimental factors.

Experimental variables spanning bioset count, host age group, viral dose and viral variants across dpi show a diversified sample set to mine for distinctive characteristics (Figure 2B-C). The 74 biosets spanned i) 5 post-infection times, ii) 3 host age groups, iii) 5 SARS-CoV-1 viral variants, and iv) 4 magnitudes of viral doses. Post-infection times were 0.5 dpi (n=6), 1 dpi (n=15), 2 dpi (n=19), 4 dpi (n=19) and 7 dpi (n=15). Age groups were divided into young mice (10 wks, n=28), adult (20 to 23 wks, n=37), and aged (52 wks, n=9). The most commonly used MA15 viral variant (n=54) induced more severe reactions in mice than MA15g (n=7) and MA15e (n=7) or ic (n=4) and TOR-2 (n=2). Intranasal infection dose levels included 100 (n=4), 1,000 (n=4), 10,000 (n=15), 22,000 (n=2) and 100,000 (n=49) viral plaque forming units (pfu). As a measure of host response, the number of significant differentially expressed genes was affected by viral variant, pfu and age of mice (Figure 2D and Supplemental Table 1). By these measures variant severity was ordered in escalation with MA15 > MA15g > MA15e > TOR-2 > ic. Two-way agglomerative hierarchical clustering of the 74 biosets showed groupings of common gene activities and sample type dispersion of host age, dpi, viral variant and viral dose (Supplemental Figure 1). Principal Component Analyses (PCA) were run to understand the influence of experimental factors, only dpi showed good separation of samples with a rotational trajectory of samples from the origin to 0.5 dpi to 1 dpi to 2 dpi and circling back toward the origin at 4 dpi to 7 dpi (Figure 2E, see portal). Figure 2F shows relative sources of variation across experimental factors, dpi was the only experimental variable to pass significance (*P* = 0.014).

### II. Discovery of Coordinated Host Response Gene Programs following Coronavirus Infection

Guided by the analysis of the 74 biosets, datasets were aggregated by dpi to generate 5 temporal meta-signatures to refine insights into the order of post-infection transcriptional events. The five temporal meta-signatures contained a total of 10,564 genes with temporal gene scores across the 5 time points split nearly even between 5,036 up-regulated genes and 5,528 down-regulated genes (Supplemental Table 2). The Venn Diagram in Figure 3A shows 410 genes common to all timepoints, whereas hundreds of genes were specific for each dpi. The highest overlap of unique genes in paired timepoints occurred between 4 dpi *vs.* 7 dpi (753 genes) and 1 dpi *vs.* 2 dpi (633 genes). The least unique overlap occurred between 0.5 dpi *vs.* 7 dpi (170 genes) followed by 0.5 dpi *vs.* 1 dpi (244 genes) and 2 dpi *vs.* 4 dpi (290 genes). These results, supported by the Principal Component Analysis in Figure 1E, indicate a clear transition between early (0.5 to 2 dpi) *vs.* late (4 to 7 dpi) infection, consistent with the early accumulation and subsequent clearance of SARS-CoV-1 or SARS-CoV-2 in mouse models (Bao et al., 2020, Glass et al., 2004), as represented in Figure 3A, left graph.

**Figure 3.**
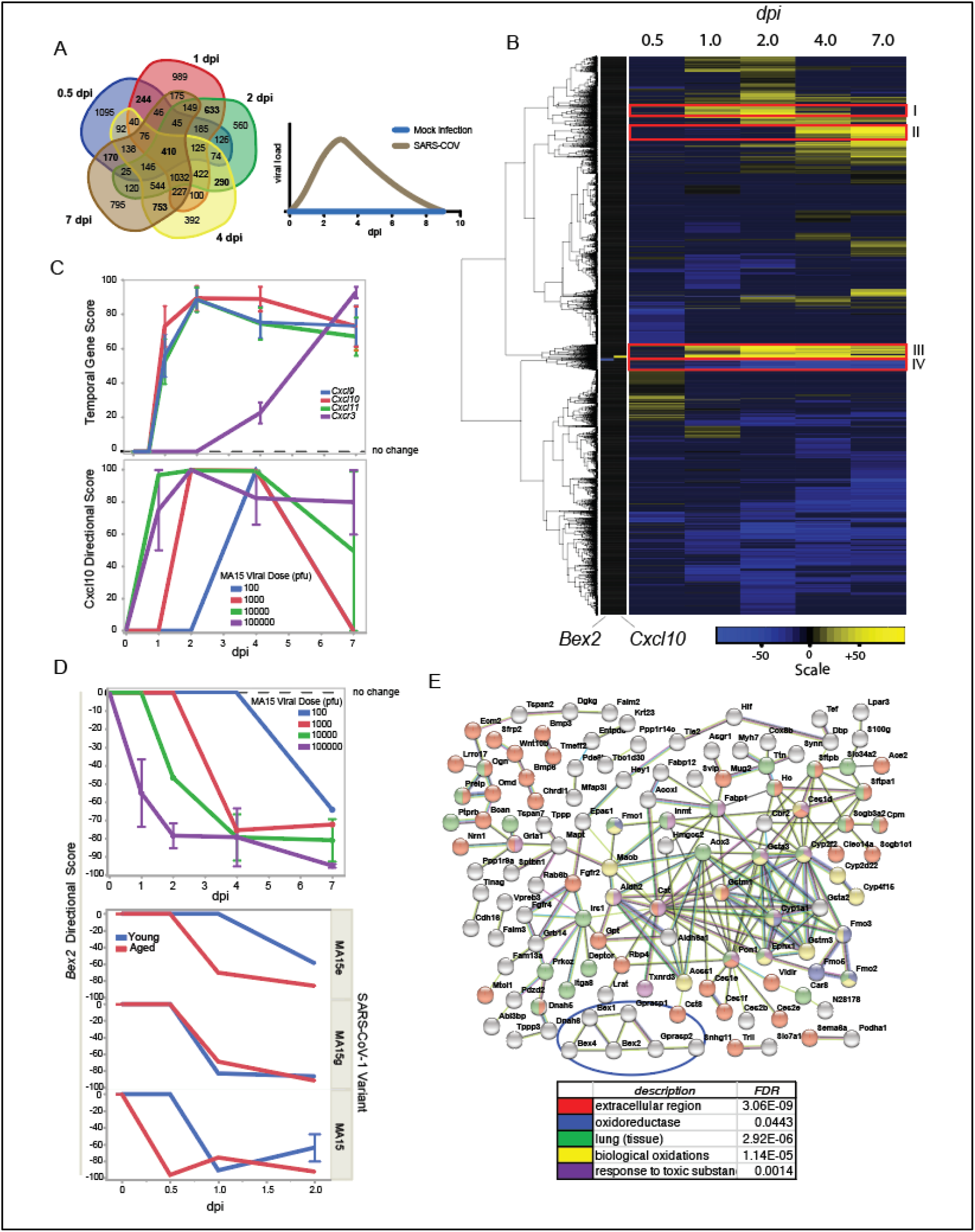
Temporal Meta-Signature Profiles Revealed Distinct Gene Programs During Coronavirus Infection. (A) The Venn Diagram shows the overlap of genes based on the temporal gene scores across the 5 dpi time points. The graph on the right shows an idealized plot of the coronavirus load *vs.* dpi from published SARS-COV-1 and SARS-COV-2 mouse models (Bao et al., 2020, Glass et al., 2004). (B) Two-way agglomerative hierarchical clustering of meta-signature genes was performed and revealed distinct gene expression patterns (I-IV), including genes that were highly up-regulated (e.g., *Cxcl10*) or down-regulated (e.g., *Bex2*) following infection. (C) The upper panel shows the temporal gene score line plots for *Cxcl* genes and the cognate receptor across dpi, whereas the lower panel shows the underlying directional gene scores by viral dose. (D) The upper panel shows the *Bex2* directional gene scores by viral dose across dpi, whereas the lower panel shows the *Bex2* directional gene scores by viral variant and age. Where adequate replicates were available mean values +/- SE are shown. (E) The 15 genes in the *Bex2* highly down-regulated cluster (pattern IV in panel (B)) were used to generate a STRING protein-protein network based on published data (see Methods).

Agglomerative hierarchical clustering revealed temporal meta-signature gene profiles that may constitute coordinated intercellular gene programs. These profiles are illustrated by the 4 patterns highlighted in Figure 3B: (I) genes that are highly up-regulated at 1-2 dpi, (II) genes highly up-regulated at 4-7 dpi, (III) genes highly up-regulated through most timepoints, and (IV) genes highly down-regulated through most timepoints. The highest-ranking up-regulated gene across all dpi (in pattern III) was *Cxcl10* (C-X-C motif chemokine ligand 10) (Supplemental Table 2). The immunoregulatory role CXCL10 has in exacerbating cytokine storm syndromes including COVID-19 has been commonly recognized (Fajgenbaum and June, 2020). The local uptake of interferon gamma by alveolar macrophages triggers release of CXCL10 that subsequently attracts a variety of lymphocytes to migrate toward and infiltrate disease tissue (Agostini et al., 2001, Coperchini et al., 2021). BSCE produced 6,059 Curated Studies containing 19,358 Biosets where *Cxcl10* dysregulation was observed across a wide range of experimental and biological designs (Supplemental Table 3). Analysis of *Cxcl10* in the BSCE Disease Atlas revealed Severe Acute Respiratory Syndrome as the most highly correlated phenotype. *Cxcl10* was also up-regulated in other top ranked infectious diseases including those caused by Adenovirus, Influenza virus and Orthopoxvirus (Supplemental Table 4). On the other hand, the highest ranked down-regulated gene (in pattern IV) across all dpi was *Bex2* (brain expressed X-linked 2), while *Bex1* and *Bex4* trended similarly downward over the course of infection (Supplemental Table 2). Genes within the *Bex* family (*Bex1*, *2*, *3*, *4*, *5*, *6*) are highly conserved in mammals and are expressed in a wide range of tissues (Alvarez et al., 2005). *Bex2* has been described as an oncogene and a dormant cancer stem cell-related protein (Kazi et al., 2015, Saijoh et al., 2021). Knockout of *Bex2* was studied in lung, where pathway analysis showed impacts on genes associated to oxygen carrier activity, PPAR signaling, immune response, and complement and coagulation cascades (Bahadar et al., 2021). Although *Bex2* remains relatively understudied in the literature to date with under 100 PubMed articles, the omic-scale data in BSCE produced 5,395 Curated Studies containing 14,278 Biosets for *Bex2* from a wide range of experimental designs (Supplemental Table 5). Severe Acute Respiratory Syndrome was the most highly correlated phenotype for *Bex2* containing biosets and was also down-regulated in other top ranked infectious diseases, including Sendai virus infection, Stachybotryotoxicosis, Legionella infection and Disease due to Paramyxoviridae (Supplemental Table 6).

Other high-ranking up-regulated chemokine transcripts included *Cxcl9* (17^th^ out of the 10,564 meta-signature gene total) and *Cxcl11* (25^th^) complementing *Cxcl10* expression (Figure 3C). These three chemotactic cytokine genes are conserved as a tandem paralog set on chromosome 5q in *Mus musculus* and chromosome 4q21 in *Homo sapiens* (Lee and Farber, 1996). They are predominately induced by interferon γ and secreted as part of a broad immune response to escalate host defenses and attract a variety of white blood cell types to sites of infection and to tumors (Callahan et al., 2021, Metzemaekers et al., 2017). The chemokine receptor CXCR3 is the cognate partner for these ligands, where binding activates pathways along the CXCL9, 10, 11/CXCR3 axis that lead to particular intracellular signaling events and systemic immune responses (Groom and Luster, 2011, Tokunaga et al., 2018). In combination, CXCL10 and CXCR3 are important for leukocyte migration to inflamed tissues (Vazirinejad et al., 2014). Temporal RNA expression of *Cxcr3* in SARS-CoV-1 infected mouse lung appears delayed with a sharp rise at 7 dpi, contrasting to 2 dpi peaks for its ligands (Figure 3C, upper panel) and coincides with infiltration of macrophages and lymphocytes late in lung infection at a time when viral titers drop in the mice (Bao et al., 2020, Glass et al., 2004). The appearance of CXCR3 at the infection site may be a biomarker for recovery.

The timing of transcriptional events was investigated by viral dose, viral variant and host age group. *Cxcl10* up-regulation shifted to earlier dpi with increasing viral dose in adult mice. (Figures 3C, lower graph). Neither severity of viral variant nor young *vs.* aged animal comparisons showed an effect on *Cxcl10* temporal scores (Supplemental Table 1). In contrast, *Bex2* expression inversely mirrored that of *Cxcl10*. Increasing viral dose was associated with an earlier onset in the suppression of *Bex2* expression, reaching maximal down-regulation at 7 dpi (Figure 3D, upper graph). In an opposite regulatory fashion, both *Cxcl10* and *Bex2* expression levels appear to be molecular read outs of viral burden (Figure 3A profile chart).

Host age and the severity of viral variant had an effect on *Bex2* expression. Down-regulation of *Bex2* occurs earlier in infection for young and aged mice treated with the most severe SARS-CoV-1 variant (MA15) compared to the weakest variant (MA15e) (Figure 3D, lower graph). The study with the highest correlation to increased expression of *Bex2* in BSCE focused on human epithelial cells exposed to hypoxic (2% O_2_) stress. In this study *Bex2* expression increased 5.6 and 4.8 fold over control levels after 12 and 48 hours, respectively (GSE136908: Tang et al, 2020). The protein-protein network software STRING (Szklarczyk et al., 2019) was used to derive deeper insight into *Bex2* and the 15 genes within the most highly down-regulated genes in the *Bex2* containing cluster IV (See Figure 3B). Though few direct relationships were observed for *Bex2*, it was connected to the G protein-coupled receptor associated sorting proteins Gprasp1 and Gprasp2. Both were strongly down-regulated across meta-signature time points (Supplemental Table 2). These genes modulate lysosomal sorting and target a variety of G-protein coupled receptors for lysosomal degradation while a lack of GPRASP2 was shown to enhance cell survival (Holmfeldt et al., 2016, Kaeffer et al., 2021). Moreover, the top phenotype in the BSCE Disease Atlas for Gprasp2 was down-regulation in Severe Acute Respiratory Syndrome [data not shown]. A highly connected hub marked by cluster IV genes *Aox3* and *Pon1* also included *Acss1*, *Aldh2*, *Aldh6a1,* Cat, *Cyp1a1*, *Cyp2f2*, *Ephx1*, *Gsta2*, *Gsta3*, *Gstm1* and *Gstm3*. All of these genes were down-regulated in the 2 dpi, 4 dpi and 7 dpi meta-signatures (Supplemental Table 2). Decreased expression of the hub genes would functionally impair their ascribed anti-oxidant and toxic chemical clearance activities. As aging leads to general loss of redox control and a well-functioning oxidative stress response, down-regulated *Bex2*, cluster IV genes and the 13 gene hub may serve as biomarkers for severe coronavirus induced decline (Chandrasekaran et al., 2017, Subramaniyan et al., 2021).

Cytokine ligands and receptors are expressed at low levels under homeostatic conditions. With cytokines such as *Cxcl10*, its activity can be induced by a variety of innate IFN-α/β and adaptive IFN-γ signals, whereas other chemokines such as *Cxcl9* are limited to adaptive IFN-γ responses (Gordon et al., 2020). Figure 4A shows the temporal scores for a variety of Interferons, Interferon Receptors, Interferon Stimulated Genes and Interferon Regulatory Factors. Interferons *Ifna1*, *Ifna2*, *Ifna4*, *Ifna12 and Ifnab* showed sharp increases in expression, peaking at 2 dpi while absent at 4 dpi and 7 dpi, whereas expression for *Ifnb1* and *Ifng* were sustained into 4 dpi and beyond. Interferon stimulated genes *Isg15* and *Isg20* had notable up-regulation spanning early and late periods. Interferon regulatory factors *Irf1* and *Irf2* were observed early while *Irf7* and *Irf9* spanned early and late periods and *Irf5* appears at 2 dpi declining into 7 dpi. Interferon receptor genes did not dramatically change across the meta-signature time course, though *Ifnar1* induction occurred early while *Ifngr2* was late. While the observed bulk tissue patterns noted here are complex, and many reminiscent of the *Cxcl10* pattern, they provide a foundation to superimpose findings with finer resolution single cell and temporal biomarker technologies.

**Figure 4.**
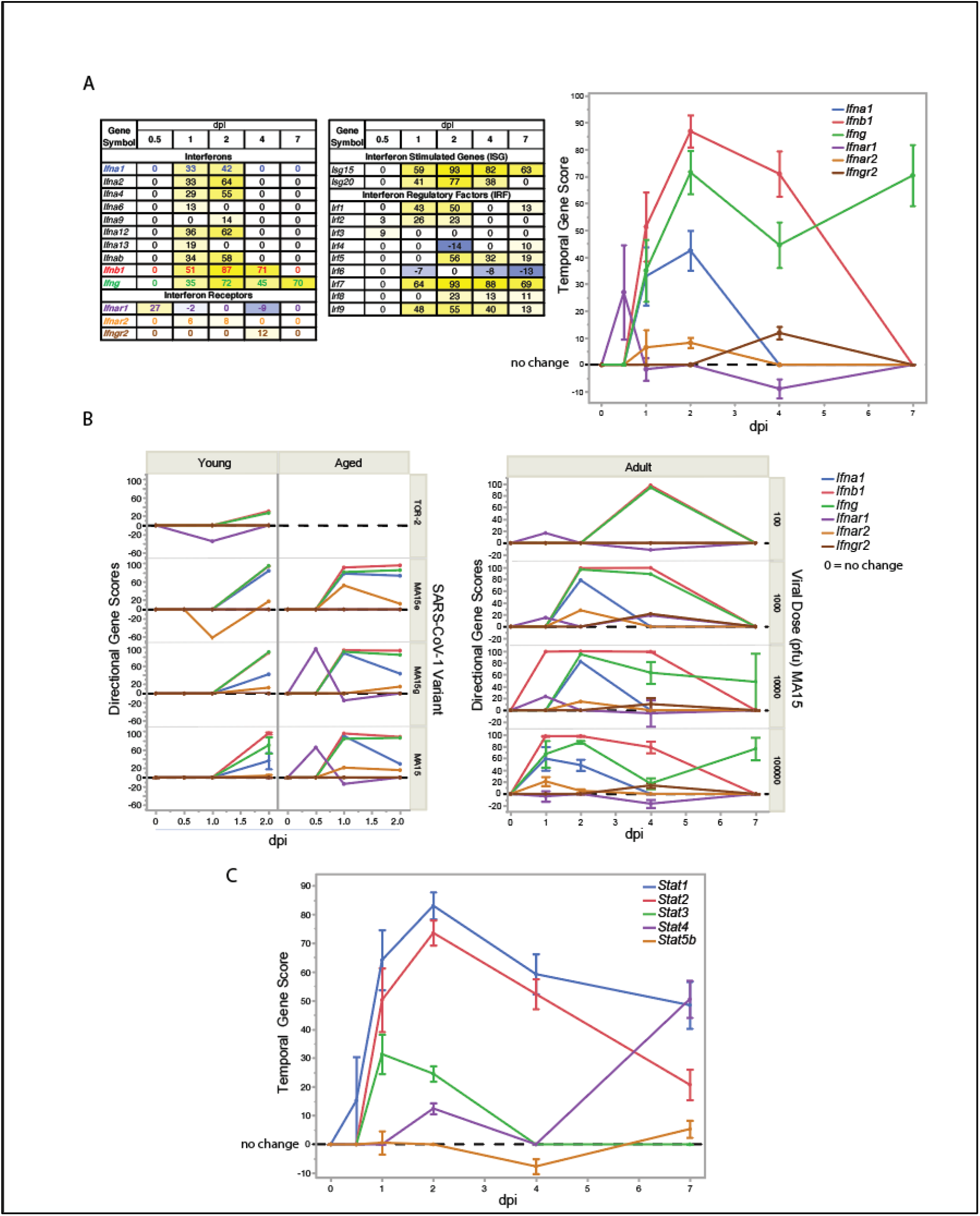
Analysis Of Potential Downstream Pathway Regulators During Coronavirus Infection. (A) The table shows a summary of the temporal gene scores for interferon related genes across dpi. The graph shows the temporal gene score profiles for selected interferon and interferon receptor genes as colored in the left table. (B) The directional gene scores underlying data in graph (A) was analyzed for young or aged mice by viral variant (left graph), and adult mice by viral dose of the MA15 variant (right graph). (C) The graph shows the temporal gene score profiles for genes of the *Stat* transcription factor family. All plots show the mean of the directional gene score and directional gene score where adequate replicates were available +/- SE.

Unlike *Cxcl10*, age effects between young and aged mice were observed for the interferon and interferon receptor genes. Peak differences in directional gene scores were seen earlier in aged mice than in young mice for *Ifna1, Ifnb1, Ifng, Ifnar1, Ifnar2 and Ifngr2* (Figure 4B, left). Data was not available for aged mice beyond 2 dpi due the lethality of the MA15 viral variant (Channappanavar et al., 2016, Roberts et al., 2007). A similar trend toward earlier maximal expression was evident for the same interferon genes and their receptors with increasing viral dosage (Figure 4B, right). The extracellular binding of cytokines to cytokine receptors commonly activates cytoplasmic STAT transcription factors in cell specific contexts (Frieman et al., 2010, Schindler and Darnell, 1995). The downstream intermediaries *Stat1* and *Stat2* showed high RNA up-regulation with temporal gene scores that peaked at 2 dpi (Figure 4C). Increasing MA15 viral dose in adult mice resulted in an earlier onset of *Stat1* and *Stat2* induction in lung, however no temporal change was observed for the MA15 variant in young *vs.* adult *vs.* aged mice (Supplemental Figure 2). In these regards, *Stat1* and *Stat2* induction patterns were similar to *Cxcl10*.

The use of the BSCE Knockdown Atlas further elucidated the relationships between immune response genes during SARS-CoV-1 lung infection. For these analyses, the 2 dpi temporal meta-signature, comprised of 4,911 genes with temporal gene scores, were correlated to published knockout models. This analysis revealed 9 knockout models of *Infar1* and 5 knockout models of *Stat1* had high negative correlations to the 2 dpi meta-signature providing further evidence for impacted gene sets (Supplemental Table 7). Studies using the knockout of *Stat1* proved to be particularly informative to interpret the host immune responses inherent in the temporal meta-signatures. One of these studied the course of SARS-CoV-1 lung infection with 100,000 pfu of the MA15 variant in *Stat1* deficient *vs.* wildtype mice (GSE36016) (Zornetzer et al., 2010). An inverse in the direction of expression, from up-regulated to down-regulated, was observed in 142 of the 148 common genes in the *Stat1* knockout bioset compared to the 2 dpi meta-signature (*P* = 6.7E-107) (Supplemental Figure 3A). In support for the relevance of *Stat1* regulatory effects, the top two functional enrichment GO biogroups in *Stat1* deficient *vs.* wildtype mice were down-regulation of both ‘innate immune response’ (*P* = 1.1E-43) and ‘response to virus’ (*P* = 2.3E-52) (Supplemental Figure 3B, C). *Cxcl10* RNA expression was the top up-regulated gene in the 2 dpi lung infected *vs.* mock infected meta-signature with an average fold change of 288 from 4 independent experiments, while in the *Stat1* knockout *vs.* wildtype bioset *Cxcl10* was reversed with down-regulation at -1.35 fold (*P* = 0.0065). While the *Stat1* knockout had no effect on *Bex2* expression, correlations with reversed expression were observed for high ranking genes across the temporal meta-signatures, including the interferon-inducible antiviral response regulators *Mx1* and *Mx2* (Haller et al., 2007, Sehgal, 2021) and other chemokines (e.g., *Cxcl11*, *Cxcl9*, *Ccl5*), interferon regulatory and stimulated genes, including *Ifit2*, *Irf7*, *Ifi44*, *Ifit1*. Reversed, anti-correlated RNA expression was observed for 2’-5’ oligoadenylate synthetase family genes (*Oas1a*, *Oas1b*, *Oas1f*, *Oas2*, *Oas3*, *Oasl1*, *Oasl2*) that trigger viral RNA degradation (Elkhateeb et al., 2016, Pulit-Penaloza et al., 2012). *Stat1* knockout reversed expression of schlafen genes (*Slfn3*, *Slfn4*, *Slfn5*, *Sfln10*) that function in cell proliferation, induction of immune responses, and regulation of viral replication (Liu et al., 2018, Mavrommatis et al., 2013, Puck et al., 2015) (Supplemental Table 7). In total, *Stat1* is shown to be a critical upstream transcriptional regulator of diverse functional pathways. In addition to the abrogation of gene expression noted, *Stat1*^-/-^ young mice infected with the most severe MA15 variant were unable to clear the infection, accompanied by elevation of neutrophils, eosinophils, and macrophages, acute respiratory distress and pulmonary fibrosis resulting in death (Frieman et al. 2010). Under the same conditions interferon *alpha* and *beta* receptor subunit 1 knockout, *Ifnar1*^-/-^, mice cleared the virus following a similar course as wildtype mice based on viral titer, weight loss, and lung immune-histochemistry results. (Zornetzer et al., 2010). Hereby, *Stat1* can be considered a central mediator of SARS-CoV-1 infectivity in mouse lung and is necessary for recovery.

### III. Functional Enrichment of Temporal Meta-Signatures

Challenges to proposing biological roles of individual genes within the SARS-CoV-1 temporal meta-signatures include the dynamic mixture of cell types and cell states in the lung samples. Functional enrichment analysis using well-structured ontological concepts provide insights to pathway level activities, which can mitigate some limitations of assigning biological processes to single genes. The 5 temporal meta-signatures were thus passed through the BSCE Pathway Enrichment application using correlations to GO biogroups (Ashburner et al. 2000), (Consortium, 2021). Exported results were aggregated into a table comprising 2,357 biogroups with directional signs and sorted by the average of the BSCE generated raw score ranks (Supplemental Table 8). The top five biogroups were ‘regulation of cytokine production’, ‘innate immune response’, ‘cell activation’, ‘inflammatory response’ and ‘response to virus’ (Figure 5A), consistent with expected biology of coronavirus infection (Perlman and Dandekar, 2005, Zhang et al., 2020). The biogroup score profiles for the top four genes in ‘Response to virus’ across the meta-signatures (*Ccl4*, *Oasl1*, *Oas2* and *Stat1*) rise in expression from 0.5 dpi to a peak at 2 dpi followed by progressively small declines between 4 dpi to 7 dpi (Supplemental Table 2). Examples of other functional gene sets perturbed during infection included correlations to lipid, peroxisome, and cell cycle processes. Subsets of ‘lipid binding’ genes show activities rising through to 2 dpi then tailing off, while other ‘lipid binding’ subsets decrease from 0.5 - 7 dpi, and the more specific child concept of ‘fatty acid metabolic process’ is distinctively more down-regulated (Figure 5B). *Acod1*, immunoresponsive gene 1, was the top ranked ‘fatty acid metabolism’ gene, it was up-regulated across all dpi (Supplemental Table 2) and may be a new target for therapeutic intervention (Weiss et al., 2018, Wu R. et al., 2020). ‘Peroxisome’ biogroup activities were also largely down-regulated by viral infection peaking early at 1 dpi and may constitute a novel target hijacked by SARS-CoV-1 (Figure 5C). *Fabp1* was the top ranked ‘peroxisome’ gene, it was second to *Bex2* as the most down-regulated gene across the temporal meta-signatures (Supplemental Table 2). A loss in host defenses due to decreased energy utilization from dysregulated fatty acid metabolism and the protection from oxidative stress afforded by *Fabp1* subverts its role in resistance to infection (Memon et al., 1999, Wang et al., 2015). Cell division concepts also show significant temporal changes. A gradual increase of up-regulated gene expression was evident in genes from ‘regulation of G1/S phase transition’ during infection. This was in stark contrast to genes in ‘regulation of G2/M phase transition’, ‘sister chromatid cohesion’ and ‘cell cycle DNA replication’, which significantly increased in expression after 2 dpi (Figure 5D).

**Figure 5.**
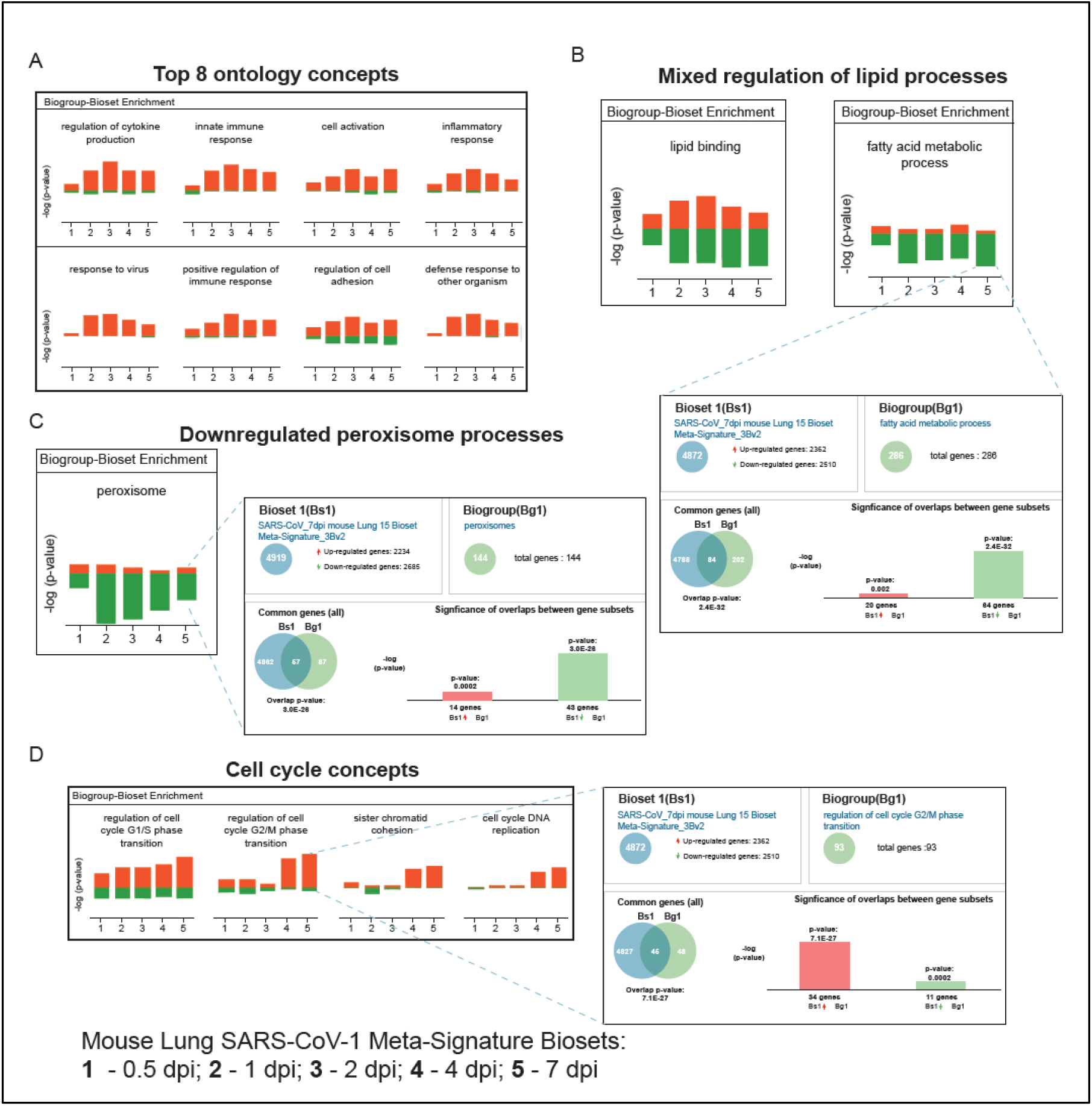
Pathway Enrichment Analyses Across SARS-CoV-1 Temporal Lung Infection Meta-signatures. The 5 temporal meta-signatures were passed through the BSCE Pathway Enrichment application using correlations to GO biogroups. Exported correlation lists were aggregated into a table comprising 2,357 concepts with directional signs and sorted by the average of the BSCE generated raw score ranks (Supplemental Table 8). (A) The panel shows the top 8 GO biogroups. (B) The panels show the mixed regulation observed between ‘lipid binding’ and ‘fatty acid metabolic process’. The blowout panel shows the correlation to the 7 dpi meta-signature. (C) The left panel shows the ‘peroxisome’ gene expression trends, whereas the blowout shows correlation to 7 dpi meta-signature. (D) The panel shows the correlations to cell cycle biogroups, while the blowout shows the correlation to the 7 dpi meta-signatures to ‘regulation of cell cycle G2/M phase transition’.

Temporal changes in cell cycle regulation were explored further given the distinctive rise of G2/M phase genes late in infection (Figure 5D). Meta-signature data was filtered by the 1,311 gene GO ‘cell cycle’ biogroup, resulting in 975 genes whose expression fluctuated over the infection time course (Supplemental Table 9). Unsupervised hierarchical clustering confirmed distinctive changes in gene expression between the early (1-2 dpi) *vs.* late (4-7 dpi) time points (Figure 6A), which suggested dynamic shifts in the regulation of the cell cycle between the peak of viral production and subsequent viral clearance (See Figure 3A). This is exemplified by the expression pattern of the proliferation markers Ki67 (*Mki67*) (Gerdes et al., 1984), *Pcna* (Kurki et al., 1986), *Ccna2* and the minichromosome maintenance complex genes *Mcm2* and *Mcm6* (Henglein et al., 1994, Stoeber et al., 2001), which peak late in infection (Figure 6B, Cell Cycle panel). This was also the pattern observed with *Ccnb1* and *Aurka, both of* which are known to define G2/M phases of the cell cycle (Fernandez and Torchia, 2019, Ryan et al., 2019). In contrast, *Ccnd2* peaked by 2 dpi, but was unchanged in the late stages. These observations help to underpin the results shown in Figure 5 and indicate an early G1/S cell cycle arrest during infection, followed by a shift to G2/M phases during viral clearance.

**Figure 6.**
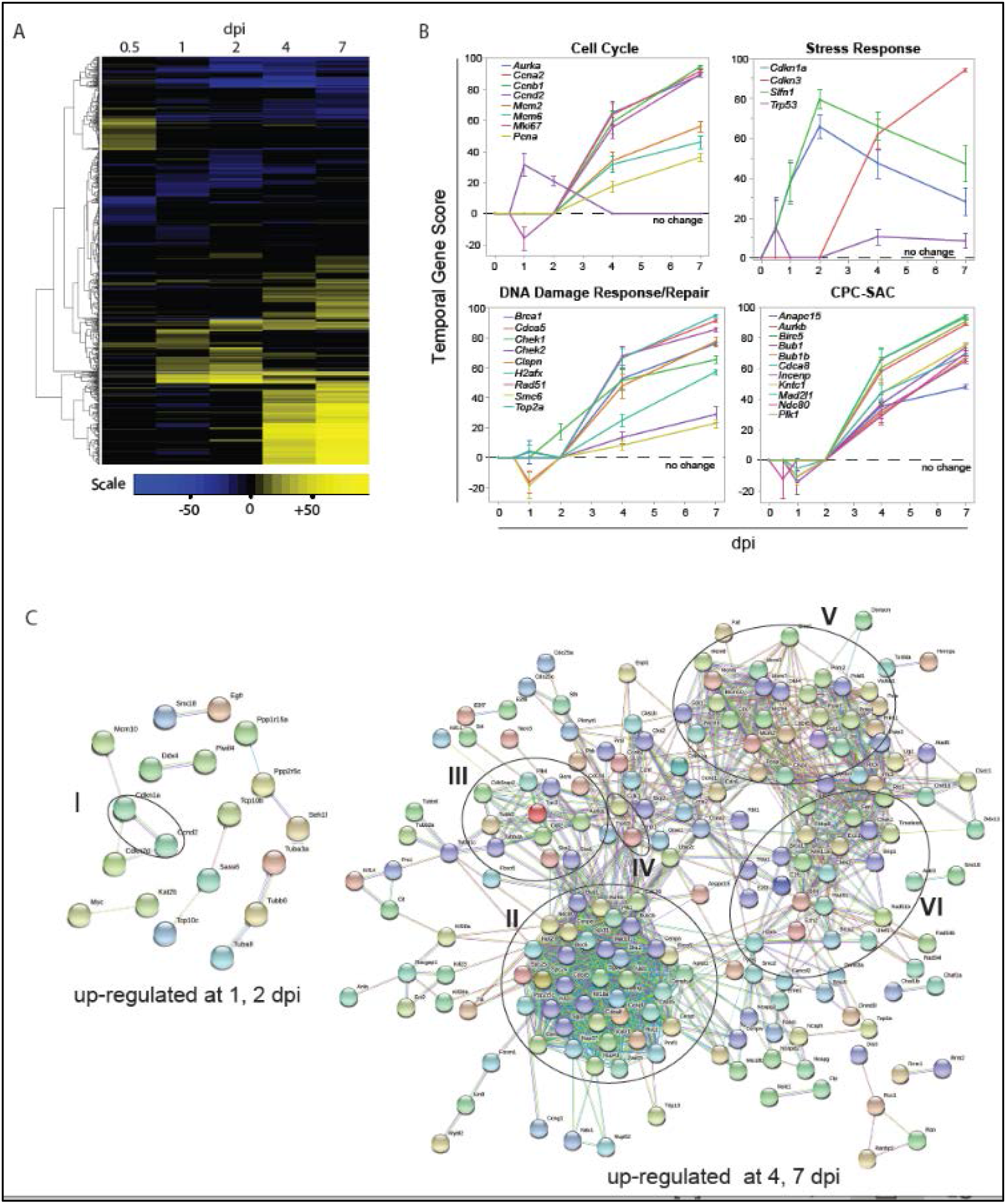
Temporal Shift In Cell Cycle Patterns During SARS-CoV-1 infection. (A) Temporal meta-signature results were correlated to the 1,311 gene GO ‘cell cycle’ biogroup using the BSCE Pathway Enrichment application. The resulting 975 genes across the 5 dpi timepoints were clustered by unsupervised methods using Euclidian distance and nearest neighbors’ agglomeration. (B) Genes from (A) that showed temporal shift were further analyzed by functional enrichment of GO and molecular pathways using WebGestalt (Liao et al., 2019, Wang et al., 2017). Distinct temporal patterns were revealed in classical proliferative markers, cellular stress response, DNA Damage/repair pathways, and chromosome attachment associated complexes (CPC-Chromosome passenger complex; SAC-Spindle assembly checkpoint). Mean values +/- SE are shown. (C) Up-regulated cell cycle genes in early (1, 2 dpi) and late infection (4, 7 dpi) were analyzed for physical complexes by the STRING protein-protein network application using high confidence evidence from the literature. Unconnected nodes are not shown. Roman numerals indicate complexes of interest (see text for discussion of hubs).

Activation of stress response pathways by genes such the Schlafen family protein *Slfn1* and the cyclin inhibitor p21 (*Cdkn1a*) may induce a G1/S cell cycle arrest. Coinciding with peak viral load, the expression of *Slfn1* and *Cdkn1a* peaked by 2 dpi but remained elevated throughout infection (Figure 6B, Stress Response panel). *Slfn1* has been shown to inhibit Cyclin D in several cell types (Brady et al., 2005, Kuang et al., 2014, Schwarz et al., 1998). The p21 protein can arrest cells at G1, S, or G2 phases in response to the activation of the cellular stress sensors such as p53 (Brugarolas et al., 1995, Radhakrishnan et al., 2004, Waga et al., 1994). The stabilization of the p53 (Trp53) protein can arrest cells or induce programmed cell death (Boutelle and Attardi, 2021). In the temporal meta-signature data, p53 expression was elevated late in infection (Figure 6B, Stress Response panel). However, p53 is continually transcribed but its protein levels are kept in check by proteolytic turnover, which under cellular stress stabilize protein levels (Boutelle and Attardi, 2021). Expression of p21 may be indicative of increased p53 activity early in infection. Lastly, the CDK2 inhibitor, *Cdkn3* also accumulated late in infection. *Cdkn3* has an essential role in the cell cycle by regulating the G1/S transition and spindle assembly check point in M-phase (Nalepa et al., 2013), however it can also affect Mdm2/p53 interactions to antagonize p21 activity (Okamoto et al., 2006). Thus, Cdkn3 may suppress or promote cellular proliferation depending on cellular context.

Analysis shown in Figure 5D revealed the enrichment of ‘cell cycle DNA replication’ and ‘sister chromatid cohesion’ events late in infection. DNA replication processes are associated with S phase and by the activation of repair pathways following DNA damage and replication stress. In the temporal meta-signature data, genes such as *Clspn, Brca1/2, and Chek1/2* (Sulli et al., 2012) (Figure 6B, DNA Damage Response/Repair panel) were up-regulated from 4-7 dpi, suggestive of the activation of DNA damage responses (DDR). While the concomitant up-regulation of genes like *Cdca5(Sororin)*, *H2afx*, *Rad51, Smc6* implicate the recovery from DNA damage (Branzei and Foiani, 2008, Schmitz et al., 2007). On the other hand, cohesion of sister chromatids is a dynamic process regulated by genes such as *Cdca5*, *Cdca8(Borealin)*, *Plk1*, *Sgo1* and *Sgo2a* (Gutiérrez-Caballero et al., 2012, Schmitz et al., 2007), which were up-regulated late in infection (Figure 6B, bottom graphs). Sister chromatid adhesion also has an important role in repair from DNA damage (Phipps and Dubrana, 2022). Following a similar pattern, kinetochore and microtubule genes were also up-regulated late in infection. These genes include the chromosome passenger complex (CPC) members (*Aurkb*, *Incep*, *Birc5*, *Cdca8*(*Borealin*), kinetochore associated genes (*Ndc80*, *Kntc1, Plk1)* and Spindle Assembly Checkpoint (SAC) associated genes (*Bub1, Bub1b, Mad2l1*) (Cheeseman, 2014, Hayward et al., 2019). Together, the complexes patrol the metaphase-anaphase transition, preventing the onset of anaphase until all chromosomes are properly bi-oriented under tension between opposing spindle poles (Hayward et al., 2019). This helps to ensure the faithful inheritance of sister chromatids to daughter cells. Persistent activation of the SAC can result in mitotic arrest. Last, there is emerging evidence that connects DDR signaling and function with that of the SAC to maintain genomic integrity (Marima et al., 2021).

STRING was used to further explore putative physical mitotic complexes between early *vs.* late infection. As shown in Figure 6C, few mitotic complexes were up-regulated and common between 1-2 dpi (Figure 6C, hub I). Notably, these complexes included *Cdkn1a* and *Ccnda*, which highlights a possible G1 cell cycle arrest early in infection. In comparison, genes common to 4 dpi and 7 dpi were suggestive of complexes associated with *Aurka* and tubulin dynamics (hub II), *Cdk1*/*CcnB1* (hub III), and the already discussed SAC signaling (IV), DNA replication (hub V), and DNA damage responses (hub VI). The number of potential cell cycle complexes was comparable between down-regulated genes early *vs.* late infection (Supplemental Figure 4). Of note, SEPTIN genes (*Sept3*, *Sept4*, *Sept8*) were down-regulated throughout infection. SEPTINS are less well understood than other cytoskeletal proteins but have roles in a variety of cell functions including anaphase of mitosis, cytokinesis, cell death, and immune system function (Woods and Gladfelter, 2021). Overall, analyses of cell cycle genes and potential complexes show a distinctive pattern of cellular stress related to viral infection.

### IV. Meta-Signature Correlations to Human COVID-19 Transcriptomes

Human COVID-19 single cell RNA sequencing (scRNAseq) studies (n=4) were imported into BSCE to assess cell type specific contributions to host infection and the extent to which the SARS-CoV-1 murine temporal meta-signatures correlate to SARS-CoV-2 infection in *Homo sapiens*. Given the attention paid above to resolving the underlying biological mechanisms of two up-regulated meta-signature clusters from Figure 3B, each was correlated to the cell type specific expression profiles from scRNAseq published data. The first cluster was the 14 gene cluster highly up-regulated throughout infection containing *Cxcl10* (Figure 3B, cluster III), it correlated highest to ‘defense response to virus’ with 10 common genes (*P* = 3.7E-25) (Supplemental Table 10). The second cluster contained 28 genes highly up-regulated at 4-7 dpi with no significant expression changes at 0.5-2 dpi (Figure 3B, cluster II), it correlated highest to ‘cell division’ with 18 common genes (*P* = 3.50E-34) (Supplemental Table 10).

The 14 genes in the *Cxcl10* cluster correlated to monocyte-macrophage lineages in human COVID-19 subjects (Table 1). Using bronchoalveolar lavage fluid from healthy, moderate, and severe COVID-19 subjects, Liao et al. found severe cases contained higher proportions of macrophages and defined four macrophage groups based on RNA expression patterns (Liao et al., 2020). The *Cxcl10* 14 gene cluster correlated highest, with 12 genes in common to IFN-γ differentiated M1-like macrophages that promote inflammation by secreting cytokines and chemokines (*P* = 3.70E-21). Results from three independent scRNAseq studies using peripheral blood mononuclear cells from COVID-19 subjects revealed the *Cxcl10* cluster correlated highest with i) 10 genes in common to CD163hi Monocytes-Cohort 1-Cluster 2 in (Schulte-Schrepping et al., 2020) (*P* = 1.90E-15), ii) 9 genes in common to the Severe COVID-19 up-regulated gene set in (Lee et al., 2020) (*P* = 2.3E-15) and iii) 6 genes in common to CD16 Monocyte-Cluster28 in (Wilk et al., 2020) (*P* = 1.70E-07). See Supplemental Table 11 for gene level details containing the correlation results. An expression matrix with *Cxcl10* cluster genes and top scRNAseq correlations to the 4 human COVID-19 studies is in Table 2. The interferon stimulated gene *ISG15* stands out as having the highest fold change in severe COVID-19 patients and is strongly represented in the 3 other COVID-19 scRNAseq studies. The SARS-CoV-1 temporal gene scores for the interferon stimulating gene *Isg15* were shown in Figure 4A. Lee et al., 2020 also observed *ISG15* commonly up-regulated in post-mortem COVID-19 lung tissue. As such, the 14 genes in the *Cxcl10* cluster may constitute a good multi-gene biomarker for early and late infection risk for severe COVID-19.

**Table 1.**
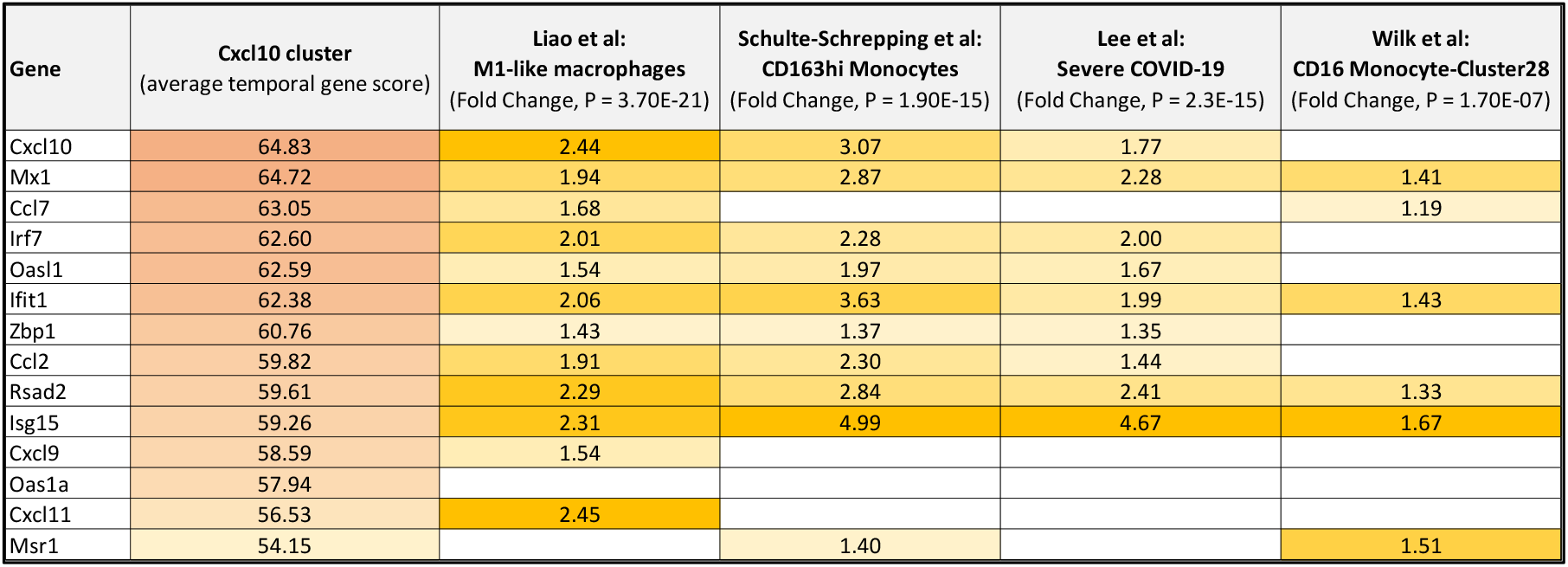
Gene Expression Matrix for the 14 Gene *Cxcl10* Up-regulated Cluster and Top single cell RNAseq Correlations to 4 human COVID-19 Studies. The values for the *Cxcl10* cluster come from the average temporal gene score (Supplemental Table 2). The values for the 4 scRNAseq studies are from the reported fold changes, while the p-values are from BSCE bioset-bioset correlations.

**Table 2.** Gene Expression Matrix with the 28 Gene 4-7 dpi Up-regulated Cluster and Top single cell RNAseq Correlations to 4 Human COVID-19 Studies. The values for the 4-7 dpi up-regulated 28 gene cluster come from the average temporal gene score (Supplemental Table 2). The values for the 4 scRNAseq studies are the reported fold changes from source articles, while the p-values are from BSCE bioset-bioset correlations.

| Gene | 4-7 dpi cluster<br>(average temporal gene score) | Liao et al:<br>Proliferating Tcell<br>(Fold Change, $P = 3.6E-49$) | Schulte-Schrepping et al:<br>Proliferating cells-Cohort 2-Cluster 12<br>(Fold Change, $P = 1.5E-27$) | Wilk et al:<br>Proliferative Lymphocytes-Cluster13<br>(Fold Change, $P = 8.6E-9$) | Lee et al:<br>Uncategorized 1_Cluster<br>15<br>(Fold Change, $P = 9.1E-22$) |
| --- | --- | --- | --- | --- | --- |
| Cenpe | 33.75 | 1.60 | 1.61 | 1.32 | 1.36 |
| Shcbp1 | 32.44 | 1.26 |  |  | 1.40 |
| Birc5 | 32.07 | 1.96 | 1.94 |  | 2.03 |
| Cdca3 | 32.06 | 1.41 |  |  |  |
| Cdk1 | 31.89 | 2.10 | 1.44 |  | 1.54 |
| Cdca8 | 31.79 | 1.54 | 1.36 |  | 1.35 |
| Pbk | 31.48 | 1.24 |  |  |  |
| Cdkn3 | 31.23 | 1.71 | 1.36 |  | 1.59 |
| Ccna2 | 31.01 | 1.81 | 1.36 | 1.23 | 1.33 |
| Aunip | 30.91 |  |  |  |  |
| Aurka | 30.74 | 1.35 |  |  |  |
| Rad51 | 30.66 |  | 1.41 |  |  |
| Ccnb1 | 30.65 | 1.61 | 1.46 |  |  |
| Plk1 | 30.18 | 1.58 | 1.48 |  |  |
| Hist1h1b | 29.90 | 1.88 |  | 1.48 | 1.65 |
| Kif20a | 29.88 |  |  |  |  |
| Spag5 | 29.69 | 1.25 | 1.58 |  |  |
| Tk1 | 29.45 | 1.84 | 1.55 | 1.23 | 2.10 |
| Cep55 | 29.43 | 1.43 | 1.35 | 1.26 |  |
| Aurkb | 29.29 | 1.61 |  |  | 1.34 |
| Sgo1 | 29.25 | 1.43 |  |  |  |
| Nuf1 | 29.18 | 1.51 |  |  | 1.28 |
| Prc1 | 28.95 | 1.31 | 1.34 |  | 1.22 |
| Cenph | 28.90 | 1.31 | 1.32 |  | 1.29 |
| Ckap2 | 28.49 | 1.38 | 1.20 |  | 1.21 |
| Melk | 28.28 | 1.28 | 1.30 |  |  |
| Asf1b | 27.80 | 1.61 | 1.65 |  | 1.37 |
| Spc25 | 27.78 | 1.44 |  |  | 1.24 |

The 4-7 dpi up-regulated 28 gene cluster was also correlated to the four COVID-19 scRNAseq studies: 25 of the 28 genes in this cluster correlated highest to those observed up-regulated in Proliferating T cells in (Liao et al., 2020) (*P* = 2.2E-36); 17 of 28 genes in Proliferating cells-Cohort 2-Cluster 12 (Schulte-Schrepping et al., 2020) (*P* = 3.2E-19); 5 of 28 genes in Proliferative Lymphocytes-Cluster13 (Wilk et al., 2020) (*P* = 7.0E-06) (Supplemental Table 12). The prevalence of up-regulated kinase genes highlights a prominent activation of intracellular signaling processes in late stage infection (*Aurka*, *Aurkb*, *Cdk1*, *Tk1*, *Plk1*, *Pbk*, *Melk*), as described previously in this article. While the observation of proliferating cells correlating to the 28 gene cluster is interesting, its impact on COVID-19 patient decline or recovery needs further study.

### V. Temporal Meta-Signature Correlations to Drug Treatments

The preceding analyses provided novel insights into the host transcriptomics and systems biology of temporal coronavirus lung infection. The analyses also described the relevance in the molecular etiology to human SARS-CoV-2 infection. Given the need for development of targeted, stage-specific treatments for COVID-19 and future coronavirus infections, the BSCE Pharmaco Atlas application was used to obtain a matrix of compound correlations to the temporal meta-signatures. This resulted in the identification of 1,706 compounds with effects on gene expression either positively or negatively correlated to the SARS-CoV-1 meta-signatures (Supplemental Table 13). Positive correlations indicate compounds have similar RNA expression activities as those induced by SARS-CoV-1 lung infection. Compounds exhibiting a negative correlation have inverse effects on post-infection gene expression activities, as in up-regulated ‘infected *vs.* uninfected’ genes are inversely observed as down-regulated in independent ‘treated *vs.* control’ biosets, thereby exhibiting potential in alleviating mechanisms gone awry in SARS (Lamb et al., 2006). Representing immunologic intervention, there were 10 Biologic - Immune Factor Antagonists and 17 Immunological Factors. Representing intracellular signaling cascades, there were 3 Biologic - Kinase Antagonists, and 66 Small Molecule Kinase Inhibitors. Other interesting Compound Classes included, Immunosuppressive drug and Anti-parasite, EZH2 inhibitors, Topoisomerase Inhibitors, DNA Methyltransferase Inhibitors, Nucleoside inhibitors, Histone Methyltransferase inhibitors, Histone Deacetylase Inhibitors, Phosphatase Inhibitors, Nonsteroidal Anti-Inflammatory Drugs, Glucocorticoids, Estrogenics, Androgenics, DNA Intercalators, Tubulin Disruptors, PPAR agonists – Fibrates, PPAR agonists - Glitazones, Statins, and Natural Products like beta-cryptoxanthin and aloe. The diversity of compound mechanisms of action reflects the breadth of viral induced dysregulated functions that evolve from early to later stages of infection.

Furthermore, spanning the broad array of mechanisms were 24 correlated compounds in Clinical Trials for COVID-19 (Table 3). Attention was drawn to the compounds with negative correlations across time. Small molecule and biologic kinase inhibitor targets included, 3 TNF antagonists, 2 JAK inhibitors as well as IL17A, IL1 receptor, SYK and BTK inhibitors. Dexamethasone had strong negative scores across all time points, while Hydroxychloroquine had strong negative scores only at 1 dpi, 2 dpi and 4 dpi that emphasizes the potential to specifically target early viral accumulation and subsequent clearance stages based on ensemble transcriptomic analyses.

**Table 3.**
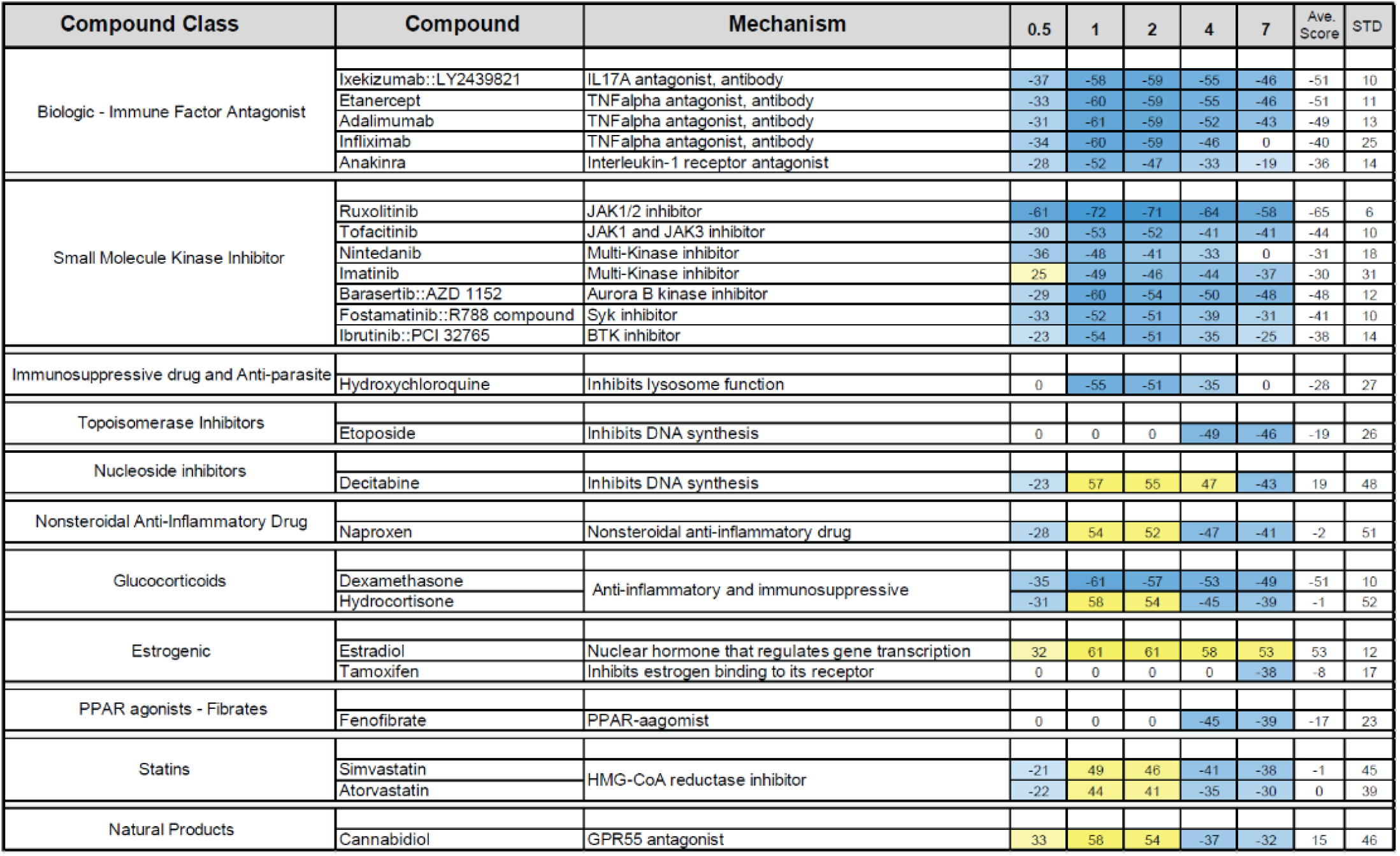
Compound Correlations to SARS-CoV-1 Lung Infected Temporal Meta-Signatures in COVID-19 Clinical Trials. BSCE Pharmaco Atlas assigns the highest ranking compound a Score of 100 in each of the 5 temporal Meta-Signatures. Lesser ranking compounds at each time point are linearly normalized to the highest ranking compound, as described in Methods. A directional sign is added to reflect a positive (yellow) or negative (blue) correlation. Compounds known to be in Clinical Trials were derived from https://ghddi-ailab.github.io/Targeting2019-nCoV/clinical. The entire list of all 1,706 significantly correlated compounds from the BSCE Pharmaco Atlas and an annotated list of other interesting compounds is in Supplemental Table 13.

### VI. Supervised Machine Learning on 74 SARS-CoV-1 Biosets and COVID-19 *in silico* Biomarker Validation

Based on the findings above, an orthogonal informatics approach was performed to define a minimal gene set that could distinguish stages of coronavirus host response using elastic-net logistic regression combined with bootstrapping (Barretina et al., 2012, Zou and Hastie, 2005). This supervised machine learning method started with 100 random splits of the 74 biosets into 2 groups based on early (0.5-2 dpi) and during the viral clearance stage of infection (4-7 dpi). Regularization was implemented to avoid overfitting because the number of sample biosets (n=74) was much smaller than the number of genes (10,564) in the dataset. The elastic-net method, which combined L1 and L2 regularization, has the flexibility in selecting the feature genes by tuning the hyperparameter “C” and “l1_ratio” (see Methods). Bootstrapping 100 times on random 80% training and 20% test bioset splits (Figure 7A) yielded 21 genes present in >90 of the bootstraps with accuracies and directional weights (Figures 7B, C). Genes were ranked by the average weights in the model. A gene with a higher negative average weight means it is relatively more important in the model in predicting the sample is at an early stage, while higher positive weights predict a sample is late stage. Six genes were predicted as early and 15 genes as late. The directional gene scores from the 74 biosets are shown in the heatmap where up-regulation dominates final assignments. A network diagram of associations based on literature assertions rendered in Pathway Studio shows connections of gene relationships spanning early stage extracellular factors (*Adm, Il1a*) with subsequent plasma membrane (*Gbp2*), cytosolic (including *Cdk1, Birc5*) and nuclear localization (including *Mik67, Top2A*) (Figure 7D, Supplemental Table 14).

**Figure 7.**
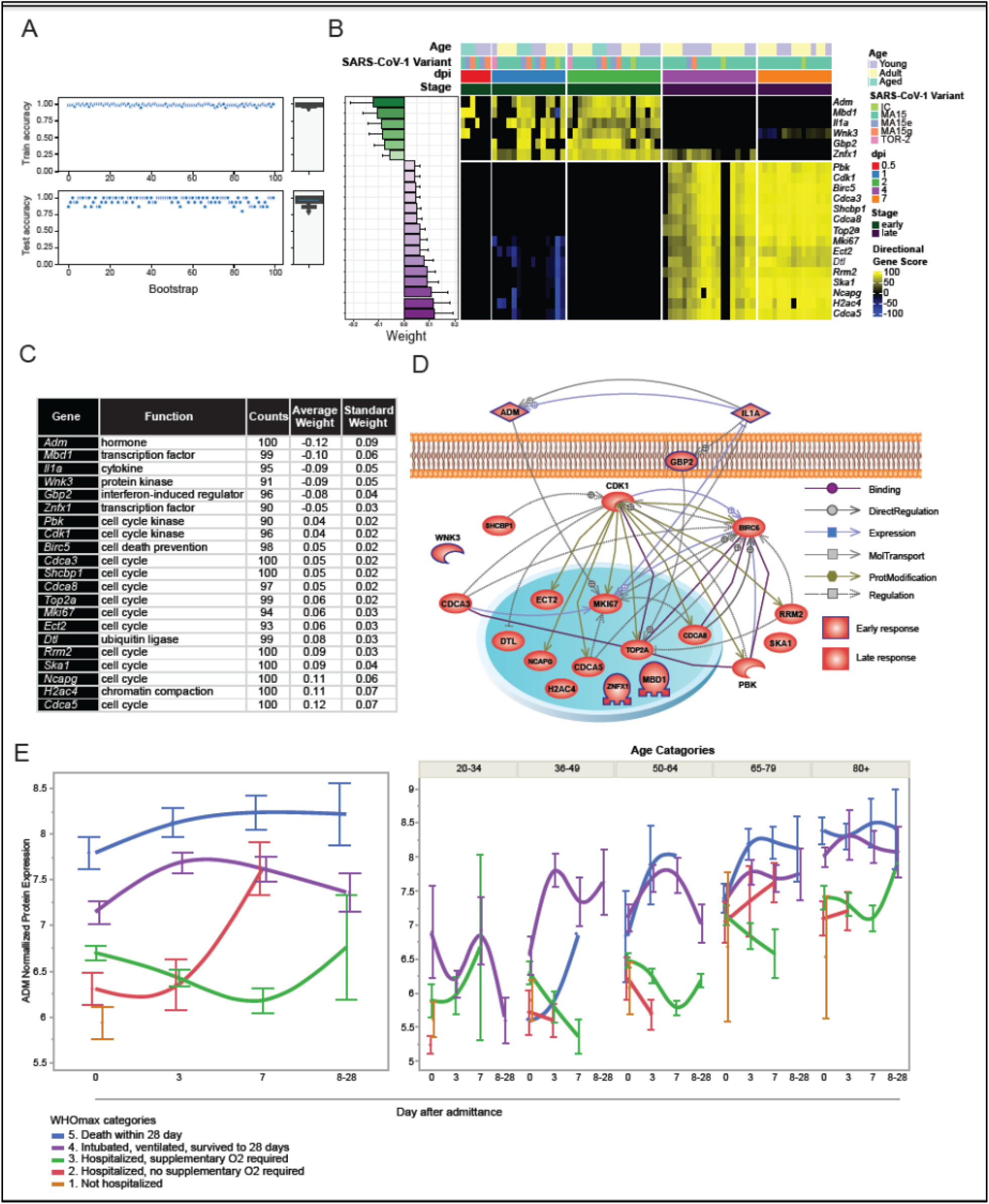
Supervised Machine Learning on 74 SARS-CoV-1 Biosets and COVID-19 *in silico* Biomarker Validation. (A) The panels show the results of training and test accuracy scores across the 100 bootstraps. (B) The heatmap shows the 21 genes present in >90 of the bootstraps horizontally ordered by dpi with the accuracies and directional weights. (C) The table lists the results with annotated gene function. Note the preponderance of cell cycle genes in late infection. (D) The relationships between the 21 gene panel was analyzed with Elsevier’s Pathway Studio. (E) The temporal data from (Filbin et al., 2021) was re-analyzed for adrenomedullin (ADM) within five profiles for escalating WHO Max categories. Mean values +/- SE are shown. The left panel shows ADM protein levels in plasma from COVID-19 humans by severity level, whereas the right panel shows ADM protein levels by severity within age groups.

Functional enrichment analyses were done on the host response stage predictor 21 gene set using the BSCE Pathway Enrichment application (Supplemental Table 15). Because there were only 6 early genes, functional biogroups were not highly populated, however ‘negative regulation of gene expression, epigenetic’ was the top GO biogroup represented by *Mdb1* and *Znfx1* (*P* = 7.7E-6). The top 2 functional enrichments for the late genes identified 8 genes associated to ‘cell division’ (p-val = 1.30E-14) and 6 genes associated to ‘chromosome segregation’: *Cdca5*, *Ncapg*, *Ska1*, *Top2a*, *Cdca8* and *Birc5* (*P* = 3.90E-12). The importance of cell cycle genes late in coronavirus infection was also evident in the analysis of temporal meta-signatures (Figures 5, 6). The most predictive late gene, *Cdca5*, cell division cycle associated 5 and also known as Sororin, was found in each of these biogroups and is understudied in the literature to date. *Cdk1* was also a late-stage predictor gene. Sororin is regulated by G2/M kinase complexes associated with *Cdk1* and *Aurkb* (Borton et al., 2016). *Cdca8* (*Boreallin*) and *Birc5* (Survivin) along with *Aurkb* are integral components of the CPC, which was discussed in the context of Figure 6B as regulating cell cycle events associated with kinetochore and microtubule dynamics late in coronavirus infection (Cheeseman, 2014, Hayward et al., 2019). It was then remarkable to find that the AURKB inhibitor Barasertib with strong negative correlations across the temporal meta-signatures and is in COVID-19 Clinical Trials (Table 3). Combined, these observations further support altered cell cycle states are a defining characteristic of SARS-CoV-1 late host responses.

*Adm,* the top early gene, encodes the circulatory hormone adrenomedullin, it has been identified as a potential biomarker in COVID-19 (Gregoriano et al., 2021, Hupf et al., 2020). This prompted the exploration of ADM activity in published human data where the Olink proteomics technology was used to analyze the plasma of COVID-19 patients. Filbin et al., 2021 divided patients into 6 categories of increasing disease progression over multiple time points with WHOmax as the patient’s most severe state. Consistent with the current study, ADM protein levels were found highly elevated, especially in patients with the highest WHOmax categories. Notably, ADM was maximally elevated in patients who died within 28 days of being admitted to hospital (Figure 7E, left). Furthermore, elevated ADM levels were associated with increasing age, as was particularly evident in patients over 65 years old and died after 28 days (Figure 7E, right).

Plasma proteomes measure a blend of systemic contributions from resident and circulating immune cells and from damaged tissues (e.g., lung, heart, kidney, brain). The strategies and results presented culminate in an unexpected discovery, where a crowdsourcing, ensemble approach using SARS-CoV-1 infected mouse lung RNA expression led to adrenomedullin, a novel protein biomarker of human SARS-CoV-2 infection and COVID-19 severity in blood.

## Discussion

### Wisdom of Crowds

Crowdsourcing of curated transcriptomic data, standard bioinformatics and advanced machine learning approaches were used to address three major questions leveraging publicly available SARS-CoV data sets: i) which genes and functional pathways are dysregulated during coronavirus infection; ii) which genes and gene sets could be used as predictive biomarkers; and iii) what compounds may counteract pathogenic processes? To address these questions, a meta-signature development approach was described to discover time dependent host infection mechanisms and shed light on potential therapeutic interventions. This approach can be adapted to any disease with sufficient RNA transcription data. However, there several limitations to the approach presented in this study: i) only RNA expression data was used to measure the biological responses and create temporal meta-signatures for host infected lung tissue. As such, translational regulation and protein activities may not correlate to RNA levels of respective genes, ii) only lung tissue was used which may not be a proxy for systemic impacts beyond the lung, iii) a mouse model with mostly unnatural SARS-CoV-1 variants may not accurately recapitulate naturally evolved causes of disease in humans and other hosts, iv) the use of only female mice across all available studies leaves open questions of mechanistic differences in males, v) BSCE used absolute fold change to rank differential expression, which minimizes genes lacking a large dynamic range vi) there are many mathematical strategies and software solutions to perform meta-analysis (Schmid et al., 2020), and is especially true in the fields of omics-scale bioinformatics and translational health science (Chang et al., 2013, Hamid et al., 2009, Ramasamy et al., 2008). Despite these limitations, the approaches used generated novel insights, some of which were corroborated by SARS-CoV-2 data. By making the data readily available and interactive, the authors encourage public efforts using it to investigate and make scientific impacts that alleviate morbidity induced by emerging SARS-CoV variants.

### Interferons, Cytokines and Stat Signaling in Host Response to Coronavirus Infection

Immune responses to coronavirus infection have taken the center stage as severe COVID-19 has been associated to an out of control ‘cytokine storm’ (Fajgenbaum and June, 2020). Under homeostatic conditions, the innate and the adaptative immune response pathways are quiescently held in wait. A complete assessment of the systemic, inter-cellular and intra-cellular factors that influence the course of immune mediated severe respiratory disease has yet to be elucidated. However, resident and infiltrating alveolar macrophages are fundamentally implicated as they produce excessive amounts of cytokines, notably CXCL10 in cytokine storm syndromes (Coperchini et al., 2021). This article provided temporal RNA expression insights to prioritize many interferon, cytokine and intermediatory factors. A 14 gene biomarker for early and late infection risk for severe COVID-19 was comprised of an assortment of immune factors. It was proposed that this multi-gene biomarker was predominantly enacted within the monocyte-macrophage axis of response to infection.

Host age was a factor whereby young animals delayed expression of key *alpha*, *beta* and *gamma* interferon and interferon receptor genes compared to adult and aging hosts. Contemporaneously, *Cxcl10* and the *Stat* family of transcriptional activators did not appear to be affected by age. However, increasing viral dose triggered *Cxcl10*, *Stat1*, *Stat2* and *Stat3* expression to appear at earlier time points lending credence as early central control factors. Furthermore, *Stat1* knockout abrogated SARS-CoV-1 induced *Cxcl10* expression (Zornetzer et al., 2010) adding support that some level of expression of these genes is necessary for recovery and in severe conditions avoiding death. Correlations of the temporal meta-signatures showed evidence that the JAK targeting JAK-STAT pathway inhibitors Ruxolitinib and Tofacitinib could effectively reverse coronavirus induced RNA expression activities from early to late stages. Extreme levels of CXCL10 measured in the blood of infected individuals of all ages may be a diagnostic marker suggesting JAK-STAT therapeutic intervention.

Lung airway microenvironments have a thin stromal framework with resident epithelial cells, T cells, B cells and other antigen presenting cells that may limit a rapid immune response and thereby enable high viral loads (Hussell and Goulding, 2010). The interplay of infected tissue resident cells signaling innate immunity and subsequent adaptive recruitment of white blood cells to transform and infiltrate infected tissues was informed by correlations of the temporal meta-signatures to COVID-19 single cell RNAseq experiments, wherein expression of the chemotactic cytokine *Cxcl10* was most highly associated to monocyte-macrophage lineages. Future spatial and single cell RNAseq studies are needed that compare bronchial lavage, tissue biopsies and blood samples from first exposure individuals to those stricken multiple times with coronavirus. After recovery from a first infection the stromal portfolio of cell types in alveoli changes to include memory B cells and T cells. Temporal perspectives on how these new sentinel cell types perform will further delineate well controlled versus out of control responses that may lead to permanent tissue damage.

### Cell Cycle in Host Response to Coronavirus Infection

Cell cycle genes showed the most dramatic changes in expression between early active viral infection and late viral clearance in the temporal meta-signature data. This pattern reflects the complicated biology of coronaviruses infecting individual cells, host viral responses, and tissue repair and return to homeostasis during viral clearance. Nevertheless, the overexpression of G2/M, DNA damage and repair response, and SAC genes late in infection have implications for long term tissue recovery from coronavirus infection. In severely affected COVID-19 individuals cytokine storm appears responsible for many of the complications and mortalities seen from SARS-CoV-2 infection after viral levels have dropped (Castelli et al., 2020, Stankiewicz Karita et al., 2022). Many of the complications associated with severe COVID-19 infections are due to multiorgan tissue damage (Mokhtari et al., 2020, Zaim et al., 2020). Dying cells are a source of growth factors by the “phoenix” pathway (Li et al., 2010), which may activate resident stem cells to proliferate after tissue damage. The induction of the phoenix pathway may then allow the switch from a G1/S cell cycle arrest early in infection when viral production peaks to a more proliferative G2/M gene expression profile during viral clearance.

Dying cells may also account for up-regulation of DNA damage and repair response and SAC genes during the viral clearance stage of infection in the meta-signature data. Previous studies have shown that dying cells release DNA fragments, so called cell free DNA (cfDNA), from the nucleus and mitochondria after severe infection or sepsis (Aucamp et al., 2018). Uptake of cfDNA by healthy cells can lead to DNA double stranded breaks and DNA damage response (Mittra et al., 2017). More important, cfDNA act as damage-associated molecular patterns that triggers inflammation (Aucamp et al., 2018). On this track, Shabrish and Mittra proposed that cfDNA from severe COVID disease may be responsible for cytokine storm and worst patient outcomes (Shabrish and Mittra, 2021). Supporting this hypothesis, serum cfDNA levels were recently correlated with inflammation, kidney tissue injury and worst disease progression and mortality in COVID-19 patients (Andargie et al., 2021). Thus, the temporal meta-signature data may, in part, reflect the effects of cfDNA during the recovery phase of coronavirus infection. It is also important to note that the DNA repair process can be error prone, leading to chromosomal abnormalities and incorporation of mutations (Phipps and Dubrana, 2022). Moreover, recent studies have begun to appreciate the role of genetic mosaicism, the heterogenous genetic changes that are present or accumulated across an individual tissue, in the progression of a variety of diseases including cancer (Hammond and Loghavi, 2021, Zekavat et al., 2021). Genetic mosaicism that happens in somatic cells can increase with age or be caused by environmental factors, and likely reflects accumulation of mutations or genetic changes in resident tissue stem cells (Behjati et al., 2014, Hammond and Loghavi, 2021, Jacobs et al., 2012, Martincorena et al., 2015). As others have noted, there may be increased predisposition for future disease due to altered immune responses after COVID-19 (Costa et al., 2022, Jafarzadeh et al., 2022, Rahimmanesh et al., 2021). It is likely then that genetic alterations after COVID-19 due to systemic cfDNA may contribute to future disease risk including cancer. Disease surveillance and DNA analysis of symptomatic COVID-19 survivors may be needed to mitigate any incurred risk.

### Adrenomedullin in Host Response to Coronavirus Infection

Adrenomedullin, encoded by the ADM gene, was identified in this article as an early biomarker of severe COVID-19 infection by applying a machine learning algorithm to the 74 mouse lung SARS-CoV-1 infected biosets. In SARS-CoV-1 infected mouse lung, *Adm* RNA was up-regulated only in the early viral accumulation period and not in subsequent clearance stages. Adrenomedullin is a peptide hormone that is ubiquitously expressed in human tissues with local and systemic vasodilating properties (Kitamura et al., 1993, Tsuruda et al., 2019). In the serum of COVID-19 patients, high ADM protein levels were detected at all WHOmax severity levels and days post hospitalization out to 28 days. Of keynote, ADM levels were progressively higher in patients with increased age and increased WHOmax level, those over 80 years old and died had the highest levels. Because it can be measured using a minimally invasive blood draw, adrenomedullin is an excellent candidate for diagnostic and treatment development. Adrenomedullin is being pursued as a predictive biomarker for a variety of diseases including cardiovascular mortality (Gar et al., 2022) and Chronic Kidney Disease (NCT05339009). Adrecizumab, a monoclonal antibody that binds adrenomedullin, attenuates oxidative stress and is a drug in clinical trials as a targeted sepsis and COVID-19 therapy (Blet et al., 2019, Deniau et al., 2021) (NCT05156671). As adrenomedullin is expressed broadly in many tissues (Asada et al., 1999), it will be interesting to know the extent to which circulating ADM derives from non-lung sources of COVID-19 subjects as other organs succumb to this syndrome.

### Bex2 in Host Response to Coronavirus Infection

This study identified many single and multi-gene RNA expression biomarkers that are up-regulated coursing across stages of coronavirus infection. It is critical that down-regulated biomarkers are not overlooked as these may be direct targets affected by viral processes overriding host defense mechanisms. The X-linked gene *Bex2* was the highest ranking down-regulated gene identified across the SARS-CoV-1 meta-signature time course, while *Bex1* and *Bex4* followed a similar trend. The BSCE omic-scale data was rich in examples for *Bex2* down-regulation in a wide variety of infectious diseases and cancers. BEX2 down-regulation induced G1 cell cycle arrest and sensitized cancer cells to pro-apoptotic agents (Naderi, 2019, Naderi et al., 2010). It is possible that BEX genes may play a similar role during coronavirus infection, namely sensitizing infected cells or their neighbors to apoptosis. This may work via viral promotion of a pro-oxidant environment interfering with redox homeostasis and antioxidant defense systems (Khomich et al., 2018).

As its gene name implies, brain expressed X-linked 2 is located on the X-chromosome and is therefore represented as two copies in females and one copy in males. There is a well-documented gender effect on severity and mortality with COVID-19 (Gebhard et al., 2020, Jin et al., 2020, Takahashi et al., 2020), with males more likely to suffer more severe symptoms with higher mortality rates than infected females. It has been suggested that this gender difference is due to the immune response, with a more robust T-cell activation correlated to a better SARS-CoV-2 infection outcome in female and young patients compared to male and old patients respectively (Takahashi et al., 2020). Given that this gene is down-regulated in a variety of viral infections, it may be that Bex2 is a protective gene that is targeted for inhibition by the invading virus. A recent study in induced pluripotent stem cells has shown that Bex2 is capable of escaping X-chromosome inactivation (Yamamoto-Shimojima et al., 2021), thereby giving females potentially two functional copies. It should also be noted that Bex1 and 2 was found to be colocalized with the olfactory marker protein (OMP) in mature olfactory receptor neurons (Koo et al., 2005), suggesting a mechanism for loss of smell in the host after viral infection.

### Therapeutics with Potential to Alleviate Coronavirus Disease Severity

Precision medicine and personalized healthcare has become a standard goal in the development of therapeutic agents where biomarkers define particular treatments to subgroups of patients experiencing the same disease diagnosis (Cook et al., 2014, Hartl et al., 2021, Langreth and Waldholz, 1999). Coronavirus infection impacts an astonishing number of normal biological processes with disease severity often dependent on the presence of comorbidities: like age, cardiovascular disease, diabetes, obesity and cancer (Bigdelou et al., 2022). Often, therapeutic strategies target more than one mechanism with potent combinations aiming for improved outcomes, as it is for COVID-19 (Crisci et al., 2020) (DeMerle et al., 2021) (Kantar er al., 2020). Several immune factor agonists, immune factor antagonists, kinase inhibitors, peroxisome proliferation agonists and histone deacetylase inhibitors were observed to favorably anti-correlate broadly across early, mid and late stages of infection, while compounds in other classes had a mix of early, mid or late effects. With access to sufficient data, it would be beneficial to study COVID-19 severity in patients taking prescription drugs that may have improved their outcome.

The cause of viral respiratory infections is not limited to members of the beta-coronavirus species (Charlton et al.). Clinical presentation of symptoms can have substantial overlap whether the causative agent is influenza A or B viruses, respiratory syncytial virus A or B, respiratory enteroviruses, rhinoviruses, respiratory adenoviruses, human metapneumovirus, parainfluenza viruses, and other coronaviruses (Kuchar et al., 2015, Petersen et al., 2020). Diagnosis of the causative agent is not possible based on reports of symptoms, yet a common physiology suggests that common molecular phenomena drive the course of events. Medicines that alleviate symptoms and improve host condition for one viral respiratory infection could improve the condition of patients presenting with a different viral respiratory infection (Singh et al., 2020).

## Conclusions

There are several areas where additional information would better inform the value of the biomarkers and transcriptional signatures presented in this study. Incorporating data on comorbidity factors and genetic susceptibility could further refine mechanistic understanding and targeted therapeutic strategies. Layering in Structural Biology studies of coronavirus gene products using 3-D modeling and docking of host ligands would help to dynamically describe events in the viral replication and host recovery phases. A whole body network model of host response gene programs would be aided by addition of molecular observations across more tissues in bulk and in single cells over the course of infection and recovery. Integrating this information with the temporal meta-signature ‘targets’ would add more dimensions and elevate opportunities for precise mechanistic intervention. As new coronaviruses will inevitably emerge, grasping the fundamental underpinnings of infection and host reactions will better position mankind to effectively blunt future disease severity and spread.

## Data and Code Availability

The data and code are also available at GitHub (https://github.com/Mark-A-Taylor/Coronavirus_Portal.git/). An interactive portal is available at http://18.222.95.219:8047.

Any additional information required to reproduce this work is available from the Corresponding Authors.

## Competing Interests

JF, MK and ZC work for Illumina, the commercial developer of BSCE. ME, ET, MT, CM, MA report no competing interests.

## Acknowledgements

The authors thank Joseph Delaney and his team of BSCE curators at Illumina.

## Author contributions

JF, ME and ET were responsible for the conception and design of the studies, performed bioinformatics analyses, biological interpretation and manuscript drafting. MK, ZC, MT, CM and MA performed bioinformatics analyses and created the interactive portal. All authors had access to and assisted in data interpretation. All authors helped to critically revise the intellectual content of the manuscript and approved the final submission.

---

**Supplemental Figure 1:**
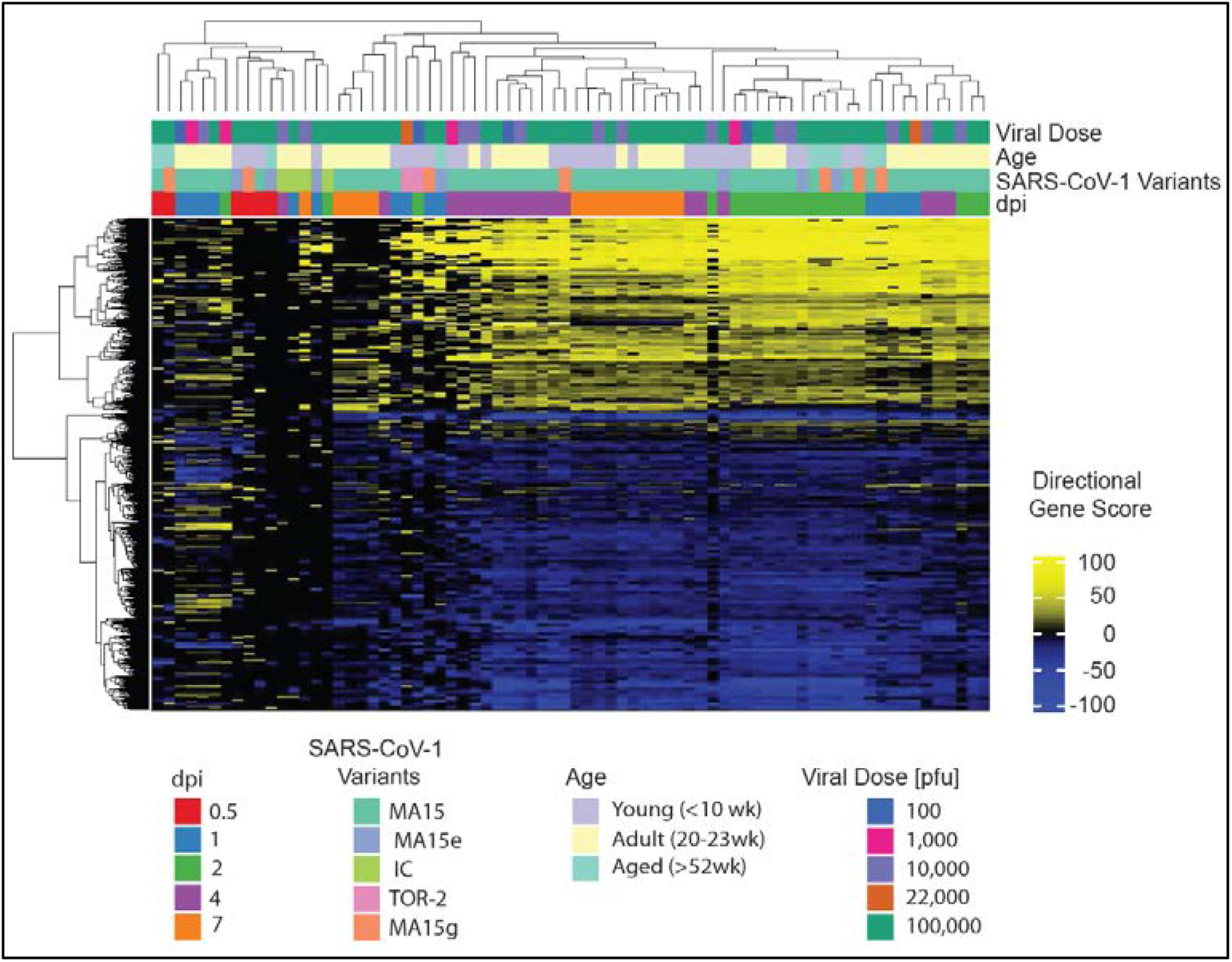
Two-way Agglomerative Hierarchical Clustering Heatmap of 74 SARS-CoV-1 Biosets. Horizontal gene and vertical sample feature orders used differential RNA Expression based directional gene scores to calculate Euclidian distance and nearest neighbors cluster agglomeration.

**Supplemental Figure 2.**
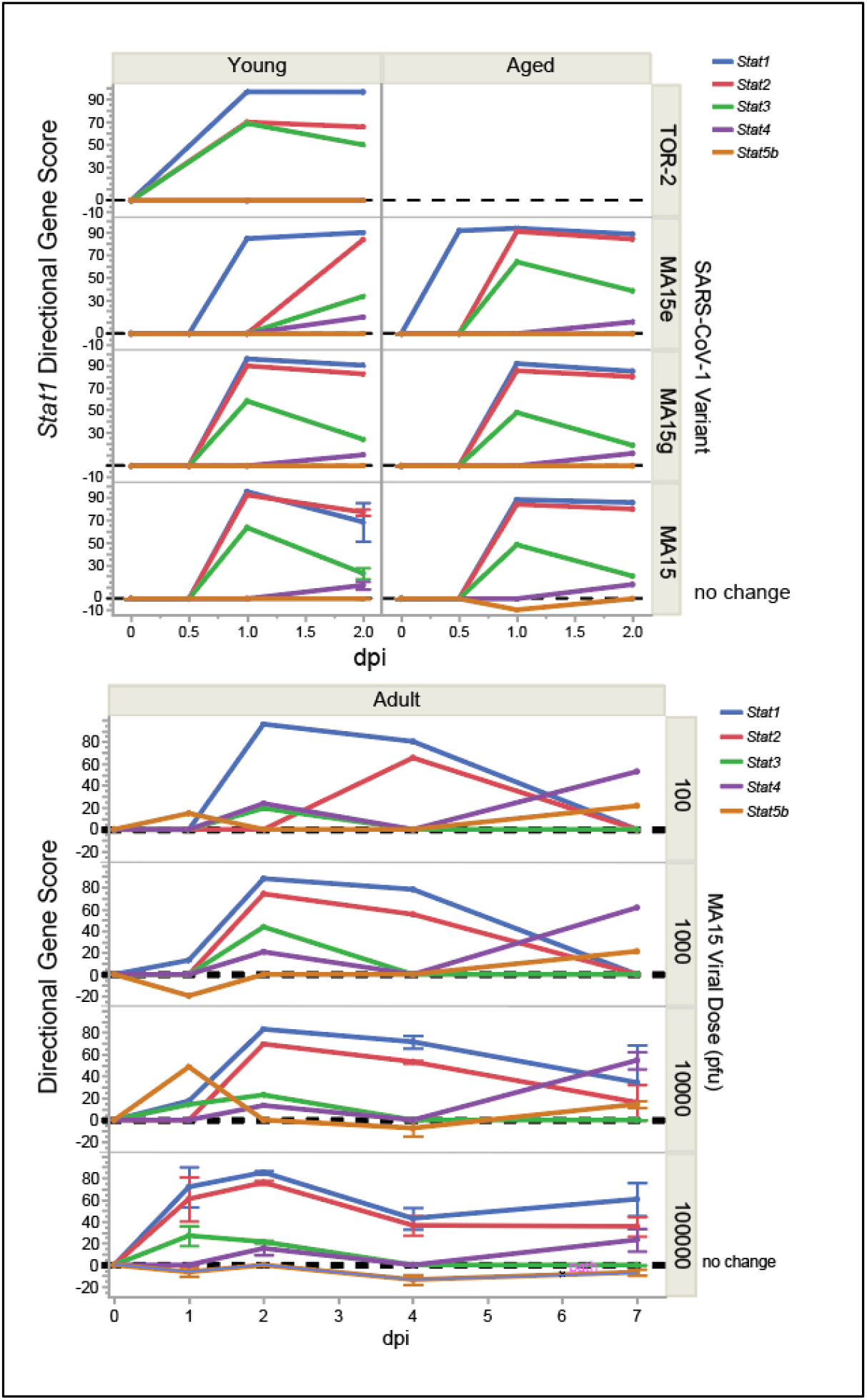
Analysis of *Stat* genes by host age, viral variant and viral dose. Top graph shows the directional gene score profiles for *Stat* gene family members in young and in aged mice by viral variants. Bottom graph shows the directional gene score profiles for *Stat* gene family member comparisons in adult mice by viral dose of the most severe SARS-CoV-1 MA15 variant. Where adequate replicates were available mean values +/- SE are shown. Note data for TOR-2 aged animals not available.

**Supplemental Figure 3.**
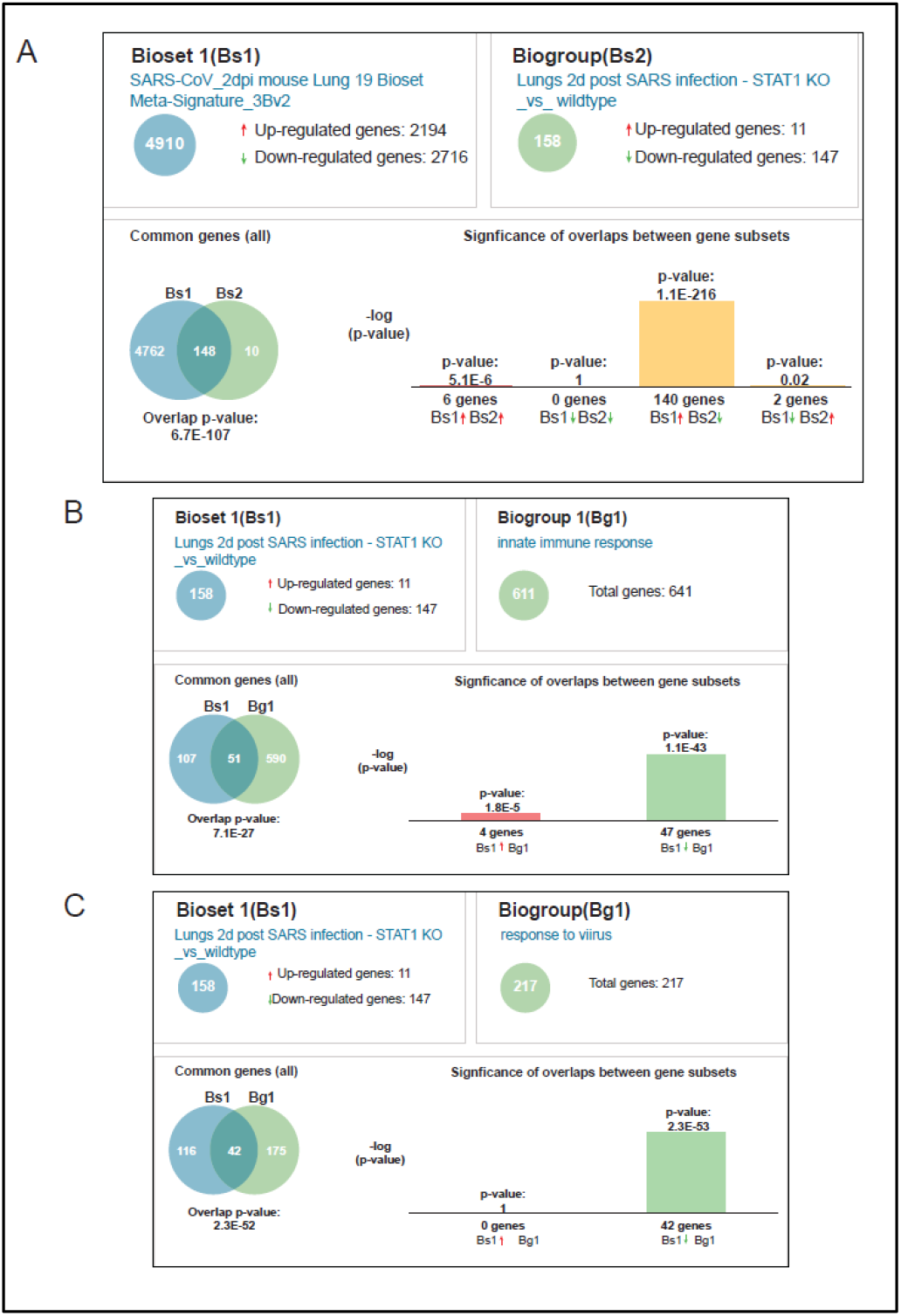
Correlation of *Stat1* Knockout Transcriptional Data to SARS-COV-1 Meta-signatures. (A) Correlations using 2 dpi meta-signature was done to compare relationships with the 2 dpi coronavirus lung infected *Stat1* knockout mouse model data from GSE36016 (Zornetzer et al., 2010). (B) The 2 dpi *Stat1* knockout bioset was compared to the GO ‘innate immune response’ genes and in (C) ‘response to virus’ genes. See Supplemental Table 7 for complete data.

**Supplemental Figure 4:**
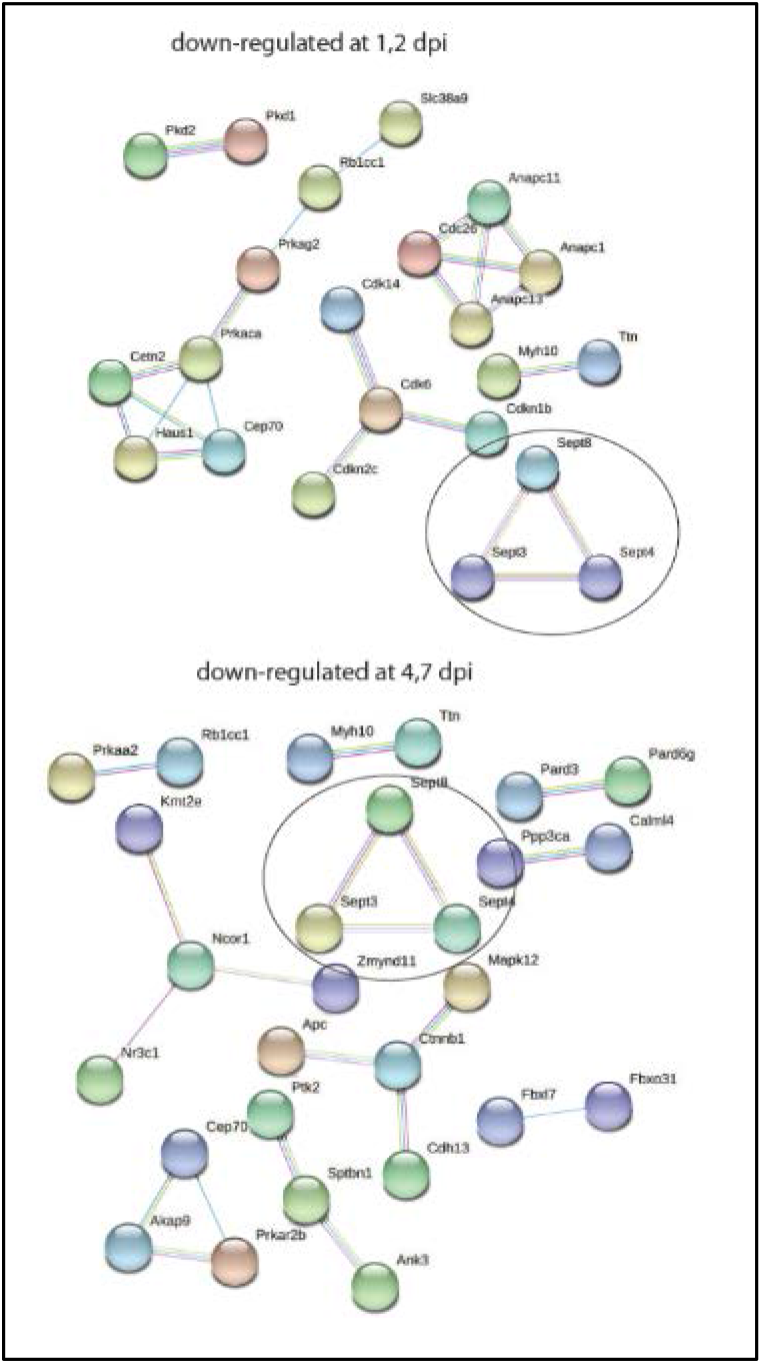
Down-regulated Cell Cycle Genes in Early (1, 2 dpi) and Late Infection (4, 7 dpi). STRING protein-protein interaction networks were generated for cell cycle genes down-regulated early and late post-infection. Network diagrams were rendered using high confidence evidence from the literature and co-expression data. Circles highlight SEPTIN genes that are down-regulated throughout infection.

**Supplemental Table 11.** *Cxcl10* 14 Gene Cluster Correlated to 4 Independent human COVID-19 single cell RNAseq Studies. Cell type specific correlations are shown divided into sections for Liao et al 2020; Schulte-Schrepping et al 2020, Lee et al 2020, and Wilk et al 2020. Cell type biosets in bold red text highlight correlations to monocyte-macrophage lineages. The bioset highlighted in bold blue text shows 9 common genes correlated a severe COVID-19 phenotype in Lee et al 2020 (*P* = 2.30E-15). P-values are from BSCE bioset-bioset correlations.

| Study Name | Bioset Name | Correlation | Common Genes | P-Value |
| --- | --- | --- | --- | --- |
| Liao et al 2020 | **Macrophage Group 1** | + | **12** | **3.70E-21** |
| Liao et al 2020 | **Macrophage Group 2** | + | **11** | **3.90E-19** |
| Liao et al 2020 | CD8pos Tcell-Moderate | + | 4 | 7.60E-07 |
| Liao et al 2020 | CD8pos Tcell-Nonexpanded | + | 4 | 1.30E-05 |
| Liao et al 2020 | CCR7pos Tcell | + | 2 | 0.0005 |
| Liao et al 2020 | CD8pos Tcell-Severe | + | 2 | 0.0012 |
| Liao et al 2020 | NK Tcell | + | 2 | 0.0023 |
| Liao et al 2020 | Proliferating Tcell | + | 2 | 0.0113 |
| Liao et al 2020 | **Macrophage Group 4** | + | **1** | **0.1182** |
| Schulte-Schrepping et al 2020 | **CD163hi Monocytes-Cohort 1-Cluster 2** | + | **10** | **1.90E-15** |
| Schulte-Schrepping et al 2020 | CD4pos T cells-Cohort 1-Cluster 11 | + | 7 | 9.00E-10 |
| Schulte-Schrepping et al 2020 | **HLA-DRlo CD163hi Monocytes-Cohort 2-Cluster 2** | + | **6** | **4.60E-09** |
| Schulte-Schrepping et al 2020 | B cells-Cohort 1-Cluster 17 | + | 6 | 1.10E-08 |
| Schulte-Schrepping et al 2020 | **HLA-DRhi CD83hi Monocytes-Cohort 2-Cluster 1** | + | **5** | **9.80E-05** |
| Schulte-Schrepping et al 2020 | Megakaryocyte-Cohort 1-Cluster 20 | + | 6 | 0.0003 |
| Schulte-Schrepping et al 2020 | NK cells-Cohort 1-Cluster 15 | + | 2 | 0.0253 |
| Schulte-Schrepping et al 2020 | **HLA-DRhi CD38hi Monocytes-Cohort 1-Cluster 1** | + | **4** | **0.0274** |
| Schulte-Schrepping et al 2020 | pDCs-Cohort 1-Cluster 8 | + | 1 | 0.0283 |
| Schulte-Schrepping et al 2020 | pDC-Cohort 2-Cluster 8 | + | 1 | 0.0297 |
| Schulte-Schrepping et al 2020 | Plasmablasts-Cohort 2-Cluster 15 | + | 2 | 0.047 |
| Schulte-Schrepping et al 2020 | Neutrophils-Cohort 1-Cluster 5 | + | 1 | 0.065 |
| Schulte-Schrepping et al 2020 | Plasmablasts-Cohort 1-Cluster 19 | + | 1 | 0.1003 |
| Schulte-Schrepping et al 2020 | **Non-classical Monocytes-Cohort 2-Cluster 4** | + | **1** | **0.2129** |
| Lee et al 2020 | **Severe COVID-19 upregulated genes** | + | **9** | **2.30E-15** |
| Lee et al 2020 | CD8+ T cell, non-EM-like_Cluster 1 | + | 2 | 0.006 |
| Lee et al 2020 | **Monocyte, nonclassical_Cluster 12** | + | **2** | **0.0311** |
| Lee et al 2020 | Dendritic cell_Cluster 18 | + | 1 | 0.0543 |
| Lee et al 2020 | **Monocyte, classical_Cluster 13** | + | **1** | **0.0751** |
| Lee et al 2020 | **Monocyte, classical_Cluster 0** | + | **1** | **0.0932** |
| Lee et al 2020 | NK cell_Cluster 3 | + | 1 | 0.1073 |
| Lee et al 2020 | NK cell_Cluster 5 | + | 1 | 0.1514 |
| Lee et al 2020 | **Monocyte, classical_Cluster 2** | + | **1** | **0.2768** |
| Wilk et al 2020 | **CD16 Monocyte-Cluster28** | + | **6** | **1.70E-07** |
| Wilk et al 2020 | Developing Neutrophil-Cluster27 | + | 6 | 6.90E-07 |
| Wilk et al 2020 | **CD14 Monocyte-Cluster3** | + | **4** | **5.00E-06** |
| Wilk et al 2020 | Neutrophil-Cluster 25 | + | 4 | 6.10E-06 |
| Wilk et al 2020 | IFN-stim CD4 T-Cluster19 | + | 2 | 0.0001 |
| Wilk et al 2020 | NK-Cluster11 | + | 2 | 0.001 |
| Wilk et al 2020 | pDC-Cluster26 | + | 1 | 0.0189 |

**Supplemental Table 12.**
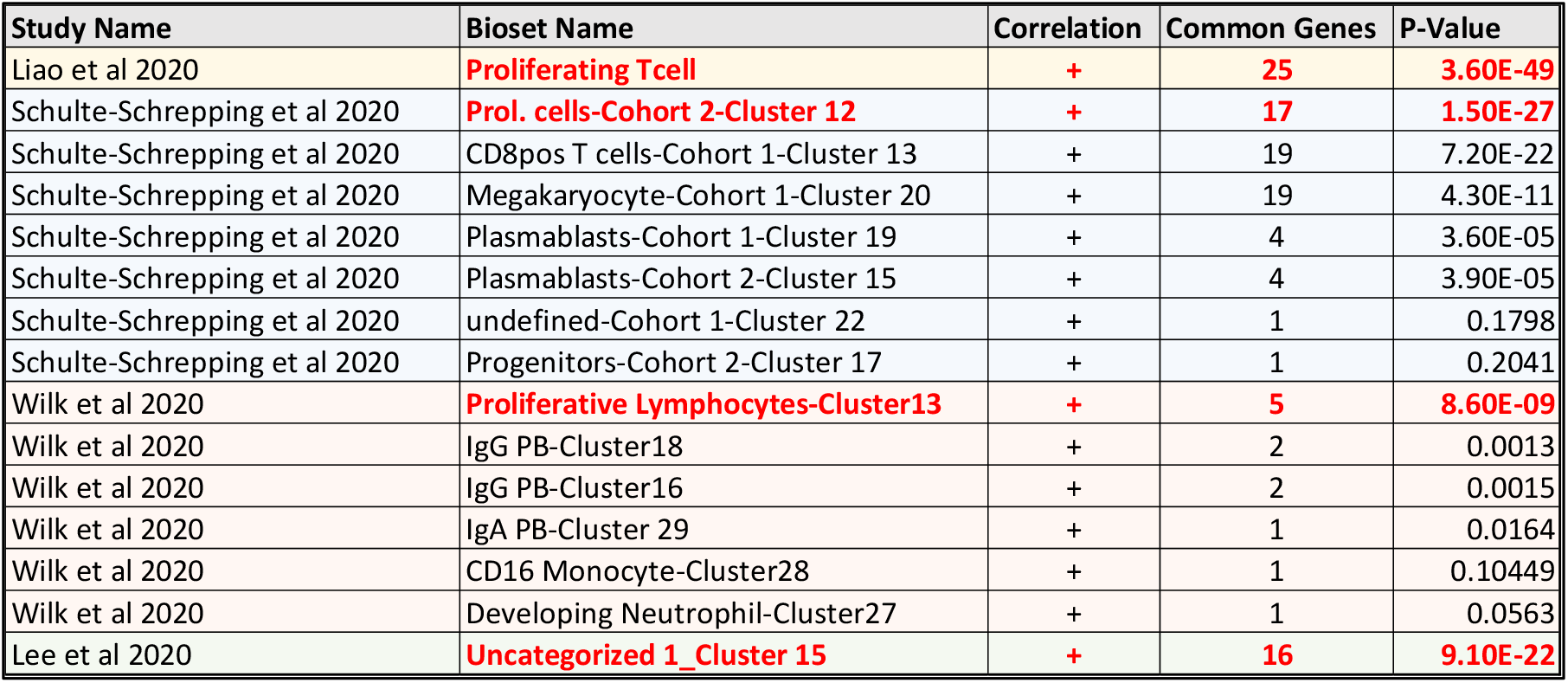
4-7 dpi Up-regulated 28 Gene Cluster Correlated to 4 independent human COVID-19 single cell RNAseq studies. Cell type specific correlations are shown divided into sections for Liao et al 2020; Schulte-Schrepping et al 2020, Lee et al 2020, and Wilk et al 2020. Cell type biosets in bold red text highlight correlations to proliferating white blood cell lineages. P-values are from BSCE bioset-bioset correlations.

## Notes

http://18.222.95.219:8047

https://github.com/Mark-A-Taylor/Coronavirus_Portal.git/

### Summary of Updates

Added Author middle initials and OrcIDs

